# 5-hydroxymethylcytosines regulate gene expression as a passive DNA demethylation resisting epigenetic mark in proliferative somatic cells

**DOI:** 10.1101/2023.09.26.559662

**Authors:** Alex Wei, Hongjie Zhang, Qi Qiu, Emily B. Fabyanic, Peng Hu, Hao Wu

## Abstract

Enzymatic erasure of DNA methylation in mammals involves iterative 5-methylcytosine (5mC) oxidation by the ten-eleven translocation (TET) family of DNA dioxygenase proteins. As the most abundant form of oxidized 5mC, the prevailing model considers 5-hydroxymethylcytosine (5hmC) as a key nexus in active DNA demethylation that can either indirectly facilitate replication-dependent depletion of 5mC by inhibiting maintenance DNA methylation machinery (UHRF1/DNMT1), or directly be iteratively oxidized to 5-formylcytosine (5fC) and 5-carboxycytosine (5caC) and restored to cytosine (C) through thymine DNA glycosylase (TDG)-mediated 5fC/5caC excision repair. In proliferative somatic cells, to what extent TET-dependent removal of 5mC entails indirect DNA demethylation via 5hmC-induced replication-dependent dilution or direct iterative conversion of 5hmC to 5fC/5caC is unclear. Here we leverage a catalytic processivity stalling variant of human TET1 (TET1.var: T1662E) to decouple the stepwise generation of 5hmC from subsequent 5fC/5caC generation, excision and repair. By using a CRISPR/dCas9-based epigenome-editing platform, we demonstrate that 5fC/5caC excision repair (by wild-type TET1, TET1.wt), but not 5hmC generation alone (by TET1.var), is requisite for robust restoration of unmodified cytosines and reversal of somatic silencing of the methylation-sensitive, germline-specific *RHOXF2B* gene promoter. Furthermore, integrated whole-genome multi-modal epigenetic sequencing reveals that hemi-hydroxymethylated CpG dyads predominantly resist replication-dependent depletion of 5mC on the opposing strand in TET1.var-expressing cells. Notably, TET1.var-mediated 5hmC generation is sufficient to induce similar levels of differential gene expression (compared to TET1.wt) without inducing major changes in unmodified cytosine profiles across the genome. Our study suggests 5hmC alone plays a limited role in driving replication-dependent DNA demethylation in the presence of functional DNMT1/UHRF1 mechanisms, but can regulate gene expression as a *bona fide* epigenetic mark in proliferative somatic cells.

## INTRODUCTION

DNA methylation is a dynamically regulated epigenetic modification with essential roles in gene regulation, genome stability, mammalian development and tissue maturation (Bird, 2002; Greenberg and Bourc’his, 2019; Li, 2002; Reik, 2007; Smith and Meissner, 2013; Wei and Wu, 2022). Mammalian DNA cytosine methylation predominantly occurs symmetrically within palindromic cytosine-guanine (CpG) dinucleotides. New CpG methylation patterns are initially established by the enzymatic activity of the *de novo* DNA methyltransferases DNMT3A and DNMT3B (Okano et al., 1999). In proliferating mammalian somatic cells, global CpG methylation patterns are maintained during DNA replication by the maintenance DNA methyltransferase DNMT1. Specifically, upon cell division, symmetrically methylated CpG patterns are inherited onto nascent DNA strands by DNMT1 and its obligate interacting partner, ubiquitin-like, containing PhD and RING finger domains 1 (UHRF1), a E3 ubiquitin-protein ligase that selectively recognizes hemi-methylated CpGs on the parental strand (Bostick et al., 2007).

Dynamic regulation of the DNA methylome involves genome-wide and locus-specific 5mC removal. Global 5mC erasure occurs in biological systems where DNA maintenance machinery is functionally impaired, precluding the re-establishment of 5mCG on nascent strands during DNA replication. For instance, DNA methylation maintenance machinery inhibition can occur when DNMT1 is excluded from nucleus (e.g. in early pre-implantation embryos) or when UHRF1 is transcriptionally repressed or functionally inhibited (e.g. in developing germ cells). Functionally, replication-dependent passive DNA demethylation is an important mechanism for erasing parental-origin specific imprints in developing germ cells, and for equalizing differences between paternal and maternal DNA methylomes in pre-implantation embryos (Greenberg and Bourc’his, 2019; Wu and Zhang, 2014).

Active erasure of CpG methylation requires 5mC oxidation by the TET family of dioxygenase proteins. TET-dependent DNA demethylation can occur through two non-mutually exclusive pathways. The first involves iterative oxidization of 5mC and 5hmC to generate highly oxidized methylcytosines: 5fC and 5caC. TDG can then selectively excise 5fC and 5caC to generate abasic sites and single-strand breaks (SSBs), which are subsequently restored to unmodified cytosines through the base excision repair (BER) pathway. By comparison to the aforementioned global DNA demethylation pathway, this TET/TDG-mediated 5fC and 5caC excision repair pathway is replication-independent and can thus operate in both proliferative and post-mitotic cells.

The second TET-dependent DNA demethylation mechanism operates indirectly, via oxidized hemi-methylation (i.e. 5hmCG/5fCG/5caCG on parental strand) induced replication-dependent depletion of 5mCG on the nascent strand. In support of this mechanism, multiple *in vitro* biochemical studies using recombinant proteins and modified duplex oligonucleotides demonstrated that DNMT1 exhibited markedly reduced activity towards the unmodified strand of hemi-hydroxymethylated CpG dyads (13-60 fold reduction compared to hemi-methylated CpG dyads) (Hashimoto et al., 2012; Valinluck and Sowers, 2007). Similarly, highly oxidized methylcytosines 5fC (8-20 fold reduction) and 5caC (>9-fold reduction) can also negatively impact DNMT1 enzymatic activity to methylate the CpG on the opposite strand (Ji et al., 2014; Seiler et al., 2018). Furthermore, UHRF1, which is essential for recruiting DNMT1 to hemi-methylated CpG dyads, has reduced affinity for selectively binding hemi-modified DNA when 5mC is replaced by 5hmC (>10-fold reduction in binding affinity for 5hmCG/CG compared to 5mCG/CG containing DNA) (Hashimoto et al., 2012). Together, these *in vitro* biochemical studies suggested a model in which oxidized forms of 5mC can substantially reduce the activity of maintenance methylation machinery (DNMT1/UHRF1) at CpG dyads thus contributing to DNA demethylation in proliferative somatic cells.

Defining the relative contributions of both pathways is critical for understanding TET-dependent DNA demethylation mechanisms. Complete inhibition of these pathways has previously been achieved by deleting all three TET methylcytosine dioxygenases, *Tet1-3*, in cultured cells (Lu et al., 2014) and in mouse embryos (Cheng et al., 2022; Dai et al., 2016). In these studies, whole-genome bisulfite sequencing analyses indicate that the loss of all TET enzymes results in hypermethylation at various *cis*-regulatory elements (CREs), including proximal promoters and distal enhancers. These findings thus support a model where TET proteins and 5mC oxidation are required for driving active DNA demethylation at dynamically regulated CREs. While 5hmC is the most abundant oxidized form of 5mC, direct evidence of 5hmC inducing replication-dependent loss of 5mC in cellular contexts remains scarce. Thus, in proliferative somatic cells, to what extent TET-dependent removal of 5mC entails indirect demethylation via 5hmC-induced replication-dependent dilution or direct conversion of 5hmC to highly oxidized 5fC/5caC for excision repair remains unclear.

In this study, we leveraged a catalytic processivity stalling variant of human TET1 (TET1.var: T1662E) to decouple the stepwise generation of 5hmC from subsequent 5fC/5caC excision and repair (**Figure 1A**). By combining locus-specific or epigenome-wide 5hmC editing (via TET1.var) with integrated multi-modal epigenetic sequencing analysis, we delineate the contributions of direct versus indirect modes of 5hmC-induced active DNA demethylation by contrasting cells expressing TET1 stalling variant (5hmC generation alone) with those expressing wild-type TET1 (5hmC generation plus 5fC/5caC excision repair). In contrast to the prevailing view, our study provides compelling evidence to support that 5hmC alone plays a limited role in driving replication-dependent DNA demethylation in the presence of functional DNA maintenance machinery DNMT1/UHRF1 but can regulate gene expression as a *bona fide* epigenetic mark in proliferative somatic cells.

**Figure 1.**
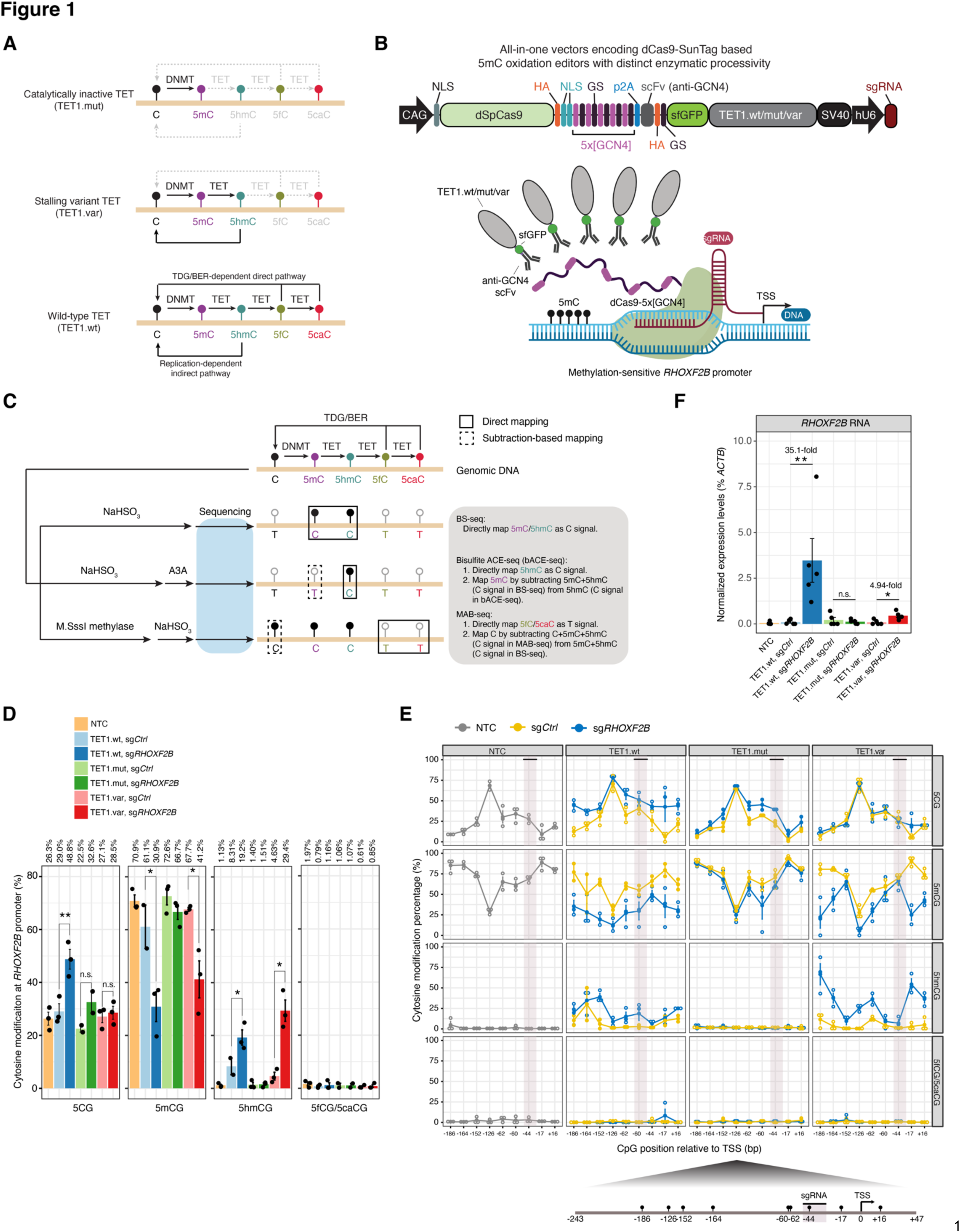
Dissecting regulatory roles of TET-mediated direct and indirect DNA demethylation pathways at specific ic locus by a CRISPR/dCas9-SunTag-based 5-hydroxymethylome editor. (**A**) Schematic overview for the impact of TET enzyme with different levels of enzymatic processivity on active DNA demethylation pathways. Wild-type TET (TET.wt) affords generation of all ox-mCs and potential to regenerate unmodified C through TDG excision of 5fC/5caC and subsequent BER. TET stalling variant (TET.var) affords 5hmC accumulation and precludes higher ox-mC generation providing a comparison against wild-type TET facilitated full active DNA demethylation to directly study 5hmC function. Catalytically inactivated TET mutant (TET.mut) identifies catalytically independent effects without editing DNA modifications. **(B)** Schematic of the All-in-one vector encoding CRISPR/dCas9-SunTag based DNA methylome editors that concomitantly express sgRNA (Top). Visual representation of the SunTag system recruiting multiple copies of human TET1 CD (wt/mut/var) to individual GCN4 peptide repeats (purple) to modify 5mC at the methylation sensitive *RHOXF2B* promoter (bottom). Abbreviations: Base Excision Repair (BER), CAG promoter (CAG), Deactivated *Streptococcus pyogenes* Cas9 (dSpCas9), Nuclear Localization Signal (NLS), General Control Transcription Factor GCN4 (GCN4), Glycine-Serine Linker (GS), single chain variable Fragment (scFv), super-folder Green Fluorescent Protein (sfGFP), human U6 promoter (hU6). The SunTag epigenome-editing schematic and experimental workflow was generated with Biorender. **(C)** Schematic depiction on the enzymatic predilections of BS/bACE/MAB-seq toward DNA modifications and their sequencing output. Integrating BS/bACE/MAB-seq affords a tetra-modal heterogenous signature that quantifies unmodified C, 5mC, 5hmC, and 5fC/5caC. Solid squares represent DNA modifications directly profiled by the assay. Dotted boxes require subtractive methods in combination with multiple assays to bifurcate modifications. Grey font delineates the enzymatic function of either sodium bisulfite (NaHSO_3_) or M.SssI Methyltransferase. Black lollipops represent DNA modifications sequenced as “C”; Empty ones represent modifications sequenced as “T”. **(D)** Locus-wide comprehensive DNA modification profiling of unmodified 5CG, 5mCG, 5hmCG, and 5fC/5caCG profiles at the *RHOXF2B* gene promoter, with modification levels (%) shown on the top. Quantification is measured by all modified CpGs as a percentage across amplicons. Individual dots represent independent biological replicates. Error bars correspond to +/− standard error. Statistical significance is established with paired one-tailed *t*-tests. *, *P*<0.05, **, *P*<0.005. **(E)** (Top) Base resolution quantities of unmodified 5CG, 5mCG, 5hmCG, and 5fC/5caCG across the *RHOXF2B* promoter [-186 to +16 bp] spanning the TSS across 9 CpG sites. Individual sites are measured as a percentage of called modification reads/total reads and further stratified by TET1 isoforms and sg*Ctrl* (yellow) versus sg*RHOXF2B* (blue). Black horizontal bar and mauve vertical bar denotes sgRNA target site. Individual open circles denote independent biological replicates. Dark circles represent mean values of each CpG site. Error bars represent +/− standard error from mean values. (Bottom) Schematic depiction of the *RHOXF2B* amplicon and individual CpG sites analyzed. Individual black lollipops represent 5mC enriched CpG sites. Relative distance to TSS is denoted by underlying numeric values. (F) *RHOXF2B* gene expression normalized against β-Actin (ACTB) expression after methylome-editing. Individual dots represent independent biological replicates. Error bars represent +/− standard error. Bar height denotes mean value amongst biological replicates. Statistical significance established with one tailed paired *t*-test. *, *P*<0.05. Non-transfection control (NTC), scrambled sgRNA (sg*Ctrl*), *RHOXF2B* promoter targeting sgRNA (sg*RHOXF2B*), non-significant (n.s). Statistical significance is established with paired one-tailed *t*-tests. *, *P*=0.0313, **, *P*=0.0245.

## RESULTS

### Development of a CRISPR/dCas9-based locus-specific 5hmC editor to investigate regulatory roles of TET-mediated direct and indirect DNA demethylation pathways

Human TET2 enzymatic processivity can be engineered to impede successive 5mC oxidation beyond 5hmC by mutating a conserved active site residue (Caldwell et al., 2021; Liu et al., 2017). Thus, we reasoned that targeted recruitment of a catalytic stalling TET variant can be a powerful approach to mechanistically characterize the direct effects of 5hmC generation on targeted genomic loci. Specifically, we introduced a threonine-to-glutamate (T1662E) mutation into the human TET1 active site (TET1.var), thus biochemically decoupling 5hmC generation from 5fC/5caC excision repair and subsequent restoration of unmodified C (**Figure 1A**).

To achieve targeted recruitment of TET1.var as a programmable 5hmC writer, we leveraged an optimized catalytically inactivated *Streptococcus pyogenes* Cas9 (dSpCas9)-based peptide repeat system (dCas9-SunTag) that enables the efficient recruitment of multiple copies of either DNMT3A– or TET1-catalytic domain (CD) for targeted DNA methylation or demethylation, respectively, with minimal off-target effects (Morita et al., 2016; Pflueger et al., 2018). We integrated TET1.var into a modular All-in-one construct that encodes all three components required for the SunTag recruitment system (Morita et al., 2016): 1) multiple short GCN4 peptide repeats are fused to a CRISPR RNA-guided dCas9 nuclease; 2) effectors (i.e. TET1 CD) are tethered to anti-GCN4 antibody single-chain variable fragments (scFv) and super-folder green fluorescent protein (sfGFP); 3) human U6 promoter (hU6) driven single guide RNA (sgRNA) (**Figure 1B**). As controls for TET1.var catalytic activity, we also constructed modular All-in-one vectors encoding wild-type TET1 (TET1.wt: capable of generating 5hmC and 5fC/5caC) and catalytically inactivated TET1 (H1671Y/D1673A TET1.mut: no 5mC oxidation activity). Immunoblotting analysis of transfected cells indicates that protein levels of both dCas9-5×[GCN4] (peptide repeats) and scFv-sfGFP-TET1 are comparable among different TET1.wt/mut/var CDs (**Figure S1A-B**).

To benchmark dCas9-SunTag based 5hmC editing efficiency, we focused on the methylation-sensitive promoter of a spermatogenesis gene, *RHOXF2B,* in HEK293T cells. Provided the relatively low expression of endogenous TETs and low global 5hmC levels observed in HEK293T cells (Ito et al., 2011), these cells afford an ideal platform for evaluating our platform. In a pilot experiment, we co-transfected HEK293T cells with dCas9-SunTag constructs (without specific sgRNAs) and a separate pU6-*sgRNA*;pEF1a-*Puro* vector encoding either scramble sgRNA (sg*Ctrl*) or an sgRNA targeting ∼200bp upstream of the *RHOXF2B* (sg*RHOXF2B*) transcriptional start site (TSS). After 24 hours of transfection, we enriched for sgRNA expressing cell populations using puromycin drug selection for 48 hours (**Figure S1A**). Next, we performed 5hmC DNA immunoprecipitation (5hmC-DIP) followed by quantitative polymerase chain reaction (qPCR) (**Figure S1C** and **Table S1**). We validated both TET1.wt and TET1.var generate substantial levels of 5hmC (TET1.wt: 16.1% and TET1.var: 17.3% of input) at the *RHOXF2B* promoter in a sgRNA– and TET1 catalytic activity-dependent manner (**Figure S1C**).

To further characterize the targeted epigenome-editing efficiency and specificity of the dCas9-SunTag system encoded by all-in-one vectors (**Figure S1D**), we performed comprehensive analysis of all major cytosine modification states of fluorescence-activated cell sorting (FACS)-isolated GFP+/DAPI-HEK293T cells to disambiguate 5mC from 5hmC (Huang et al., 2010) and unmodified C from 5fC/5caC (Wu et al., 2014). First, to resolve base ambiguity between 5mC and 5hmC, we employed BS-seq in conjunction with bisulfite-assisted APOBEC-Coupled-Epigenetic-sequencing (bACE-seq) (Fabyanic et al., 2023). Specifically, bACE-seq quantitatively profiles 5hmC at base resolution by harnessing the differential deaminase activity of human APOBEC3A (A3A) deaminase towards 5mC and chemically protected 5hmC (**Figure 1C**). Through subtracting bACE-seq from BS-seq signals, we can reveal true 5mC levels. Second, to resolve epigenetic ambiguity between unmodified C and 5fC/5caC levels, we utilized M.SssI-assisted Bisulfite sequencing (MAB-seq) (Wu et al., 2014). Together, with sample-specific lambda phage DNA spike-ins as internal controls for validating A3A deaminase efficiency (n=7, mean±sd: 98.7±0.53% 5mCG deamination rate for bACE-seq) and M.SssI methyltransferase (n=7, mean±sd: 98.0±0.20% 5CG protection rate for MAB-seq) enzymatic activity (**Figure S1E-F**), the integrated BS/bACE/MAB-seq assays afford precise, base-resolution analysis for 5mCG, 5hmCG, 5fCG/5caCG, and unmodified 5CG levels.

Because 5hmC (<0.5%) and true 5mC (<1.6%) levels were low at non-CG contexts (CHG and CHH) across samples (**Figure S1G**), we focused our locus-specific epigenetic sequencing analysis on CpG sites (9 CpGs within 290 bp amplicons) at the *RHOXF2B* proximal promoter region. Locus-wide analysis of CG modification levels reveal that in contrast to non-transfection control (NTC: 1.13%) or TET1.mut (sg*Ctrl*: 1.40%, sg*RHOXF2B*: 1.51%), targeted recruitment of both TET.wt and TET.var to the *RHOXF2B* proximal promoter markedly increases 5hmC levels (TET1.wt, sg*Ctrl*: 8.31%, sg*RHOXF2B*: 19.2%; TET1.var, sg*Ctrl*: 4.63%, sg*RHOXF2B*: 29.4%;) (**Figure 1D**). Expectedly, we also observe a corresponding decrease in 5mCG levels when TET1.wt and TET1.var are targeted by sg*RHOXF2B* to the promoter region (**Figure 1D**). While we did not observe any substantial increase in steady state 5fCG/5caCG levels across samples, unmodified 5CG levels significantly increased in TET1.wt (sg*Ctrl*: 29.0%, sg*RHOXF2B*: 48.8%) but not in TET.var (sg*Ctrl*: 27.1%, sg*RHOXF2B*: 28.5%) (**Figure 1D**). Base-resolution analysis of all 9 CpGs covered by the amplicon further confirm these observations and reveal epigenetic heterogeneity between CpG sites in response to targeted epigenome-editing (**Figure 1E**). Together, these results support a model in which highly oxidized 5fC and 5caC (generated only by TET1.wt) are rapidly removed by endogenous TDG enzyme and restored to unmodified C through excision repair in our system. As targeted recruitment of TET1.var only results in marked increases in 5hmCG but not 5CG levels, these results validated that the catalytic processivity of TET1.var is distinct from that of TET1.wt. Thus, when paired with the dCas9-SunTag system, the TET1 stalling variant can serve as robust 5hmC editor at targeted loci.

To determine the potentially divergent gene regulatory effects of TET1.wt (5hmC generation plus 5CG restoration) and TET1.var (5hmC generation alone) on the *RHOXF2B* gene expression, we performed qRT-PCR analysis of FACS-isolated GFP+/DAPI-HEK293T cells transfected with all-in-one dCas9-SunTag-TET1.wt/mut/var effectors for 72 hours (**Figure S1A**). While targeted recruitment of both TET1.wt (35.1-fold for sgRHOXF2B vs. sgCtrl, *P*=0.0245) and TET1.var (4.94-fold for sgRHOXF2B vs. sgCtrl, *P*=0.0313) results in substantial reactivation for *RHOXF2B* gene expression compared to scrambled sgRNA controls, TET1.wt was significantly more robust in transcriptional activation (sg*RHOXF2B*: *P*=0.0344 for TET1.wt vs. TET1.var) (**Figure 1F**). Given the relatively commensurate expression levels of dCas9-SunTag systems (**Figures S1B and S1D**), this difference between TET1.wt and TET1.var in the transcriptional reversal of somatic silencing of the methylation-sensitive *RHOXF2B* promoter may be predominantly attributed to distinctions in the catalytic processivity of these TET variants.

Taken together, these results suggest that dCas9-SunTag-TET1.var can efficiently oxidize 5mC to 5hmC on hyper-methylated chromatin substrates in cells, but cannot generate 5fC/5caC and restore unmodified C. Therefore, this dCas9-SunTag-TET1.var platform can serve as a versatile tool to characterize gene regulatory roles of 5hmCG at directly targeted *cis*-regulatory elements. In addition, gene expression analysis shows that 5hmC alone may have a moderate gene activating role at methylation-sensitive promoters independent of its role as an intermediate of the TET-mediated direct DNA demethylation pathway.

### Impact of the TET1 stalling variant on (hydroxy)methylomes and restoration of unmodified C across diverse genomic features

Having validated the catalytic processivity of 5hmC stalling variant (TET1.var) in the context of targeted epigenome-editing of an endogenous genomic locus, we next sought to comprehensively analyze all three major cytosine states (5C/5mC/5hmC in the CG context) in HEK293T cells that globally express TET1.wt/mut/var CD (**Figure 2A** and **S2A**). Western blot analysis indicates that the protein levels of different TET1 CDs are commensurately translated across biological replicates in HEK293T cells (**Figure S2B**). Transcriptome-wide analysis shows that while all three endogenous TET enzymes (TET1-3) are expressed at low levels, the normalized RNA levels of TET1.wt/mut/var were markedly higher (on average 38-fold) than that of endogenous TET1 in controls (**Figure S2C**). Therefore, these results suggest that 5mC oxidation activity measured in TET1.wt-and TET1.var-expressing cells is predominantly derived from exogenous TET enzymes. In addition, global expression of TET1.wt/mut/var did not detectably affect the expression levels of endogenous genes encoding all major components involved in the DNA methylation and demethylation enzymatic cascades (**Figure S2C**) (Wei and Wu, 2022).

**Figure 2.**
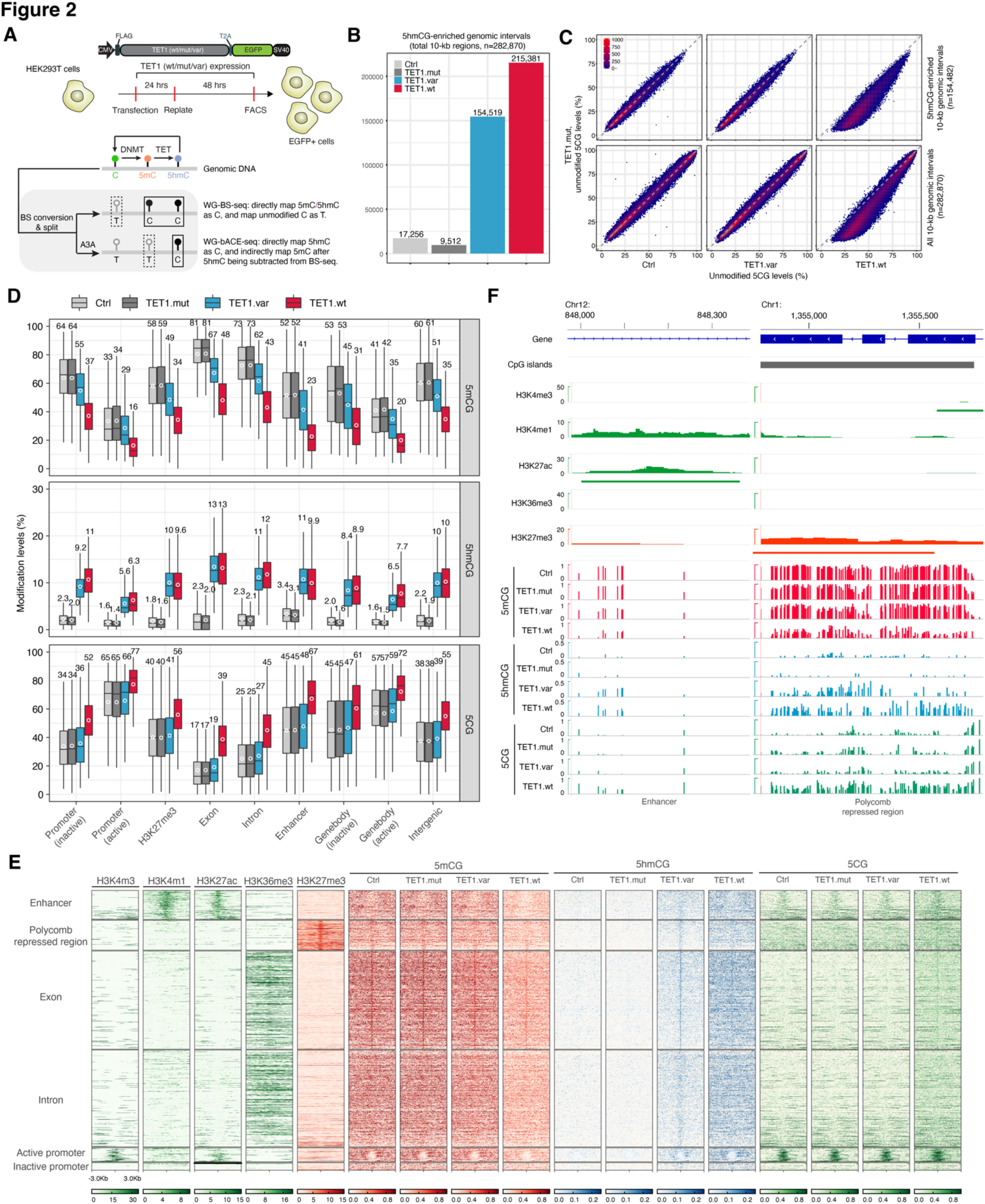
Genomic regions enriched with 5hmCG alone are not associated with global restoration of 5CG in rating somatic cells. (**A**) (Top) Schematic depiction of the over-expression vectors used for transfections experiments. hTET1 catalytic domains were generated to express either TET1 wild-type (wt), catalytic mutant (mut), or 5hmC-stalling variant (var). EGFP is used to isolate plasmid expressing cells with FACS. FLAG-tag was used to probe for protein expression. TET1 isoform over expression plasmids are transfected into HEK293T cells for 72 hours prior to isolating for GFP+/DAPI-cell populations. (Bottom) Schematic for the sequencing output for integrated whole genome (WG)-BS-seq and WG-bACE-seq to quantify 5mC, 5hmC, and unmodified C. Stagnated rectangle boxes denote enzymatic deamination by sodium bisulfite and APOBEC3A (A3A) for unmodified C and 5mCG in BS-seq and bACE-seq respectively that are read as thymine (T). Solid rectangle boxes designate DNA modifications read as cytosine (C) through sequencing output. Notably, because steady state 5fC/5caC are nearly undetectable, they have been grouped as unmodified C (C). **(B)** Bar graph of 10-kb genomic regions enriched for statistically significant level of 5hmCG (*P* value = 2.5 x 10^-4^) in control and TET1.mut/var/wt-expressing HEK293T cells, with number of 5hmCG-enriched genomic intervals listed above each bar. **(C)** Correlation density plots of 5CG levels between TET1.mut and Ctrl/TET1.var/TET.wt cells. Correlation analysis is performed with 5hmCG-enriched (top: *n* = 154,482, common to both TET1.wt and TET1.var) or all (bottom: *n* = 282,870) 10-kb genomic bins spanning the human genome. **(D)** Box plots of 5mCG, 5hmCG and 5CG levels at subsets of annotated genomic or regulatory regions enriched for 5hmCG in both TET1.var and TET1.wt cells. **(E)** Heat map representation of normalized ChIP-seq signals of major histone modifications (H3K4me3, H3K4me1, H3K27ac, H3K36me3 and H3K27me3 in wild-type HEK293T), and 5hmCG (bACE-seq), 5mCG (derived from BS-seq and bACE-seq), and 5CG (derived from BS-seq) in Ctrl and TET1.mut/var/wt HEK293T cells across a subset of annotated genomic or regulatory regions enriched for 5hmCG in both TET1.var and TET1.wt cells. The genomic features are ranked by 5hmCG levels in TET1.var cells. **(F)** Genomic track view of major histone modifications and base-resolution 5mCG (red), 5hmCG (blue) and 5CG (green) maps at two representative loci (left: H3K4me1/H3K27ac marked intragenic enhancer; right: H3K27me3-marked CG-rich Polycomb repressed regions). Only CGs covered by at least two reads are shown. BS-seq and bACE-seq tracks represent merged data sets from two biological replicates.

To jointly profile 5mCG and 5hmCG from the same sample at base-resolution, we applied an integrated (hydroxy)methylome sequencing approach by combining BS-seq (maps 5mC+5hmC) with bACE-seq (maps 5hmC only) to analyze HEK293T cells that globally express TET1.wt/mut/var CDs (**Table S2**) (Fabyanic et al., 2023). After 72 hours of transient transfection in HEK293T cells, genomic DNA (gDNA) was co-purified with total RNAs from FACS-isolated GFP+/DAPI-cells and spiked with 0.5% lambda phage gDNA as internal controls to access the sample-specific A3A performance in bACE-seq (average A3A deamination efficiency: 99.1%, benchmarked by *in vitro* methylated lambda gDNA) (**Figure S3A**). Next, we quantitatively determined global changes in 5mCG/5hmCG/5CG levels in response to TET variants with different catalytic processivity (i.e. TET1.wt/mut/var). Global expression of TET1.var (5hmCG: 3.95%) results in substantially higher level of 5hmCG levels compared to catalytically inactive TET1.mut (5hmCG: 0.92%), and TET1.var is less active than TET1.wt (5hmCG: 10.5%) in terms of 5hmC generation (**Figure S3B**), which is consistent with previous results biochemically characterizing human TET2 variants *in vitro* (Liu et al., 2017),

To gain a genome-wide view of 5hmCG levels, we analyzed 10-kb non-overlapping genomic intervals in TET1.wt/mut/var expressing cells. Specifically, paired analysis of the same pool of bisulfite converted gDNA shows that 5mCG and 5hmCG profiles are highly concordant between biological replicates (**Figure S3C**). We then merged the two replicates and performed statistical calling of 5hmCG enriched 10-kb genomic regions (*P* < 2.5 x 10^−4^) using a binomial distribution model previously established for identifying 5hmC-modified CpG sites in mammalian genomes (Schutsky et al., 2018). Compared to controls (n=17,256) and TET1.mut (n=9,512), we identified substantially higher numbers of 10-kb genomic regions that are significantly enriched for 5hmCG in TET1.var (n=154,159) and TET1.wt (n=215,381) samples (**Figure 2B**). Surprisingly, correlational analysis of unmodified 5CG levels for either all (n=282,870, top in **Figure 3C**) or 5hmCG-eriched (n=154,482, bottom in **Figure 3C**) 10-kb genomic intervals reveal that only TET1.wt but not TET1.var induces marked increases in 5CG levels when compared to TET1.mut control. Quantitative analysis of genomic intervals associated with different levels of 5hmCG show that 5CG levels only substantially increased in TET1.wt-, but not in TET1.var-expressing cells when compared to TET1.mut (**Figure S3D**). Taken together, these results suggest in contrast to TET1.wt, 5hmCG generation alone by TET1.var is not sufficient to induce substantial genome-wide restoration of unmodified cytosines (i.e. 5CG).

**Figure 3.**
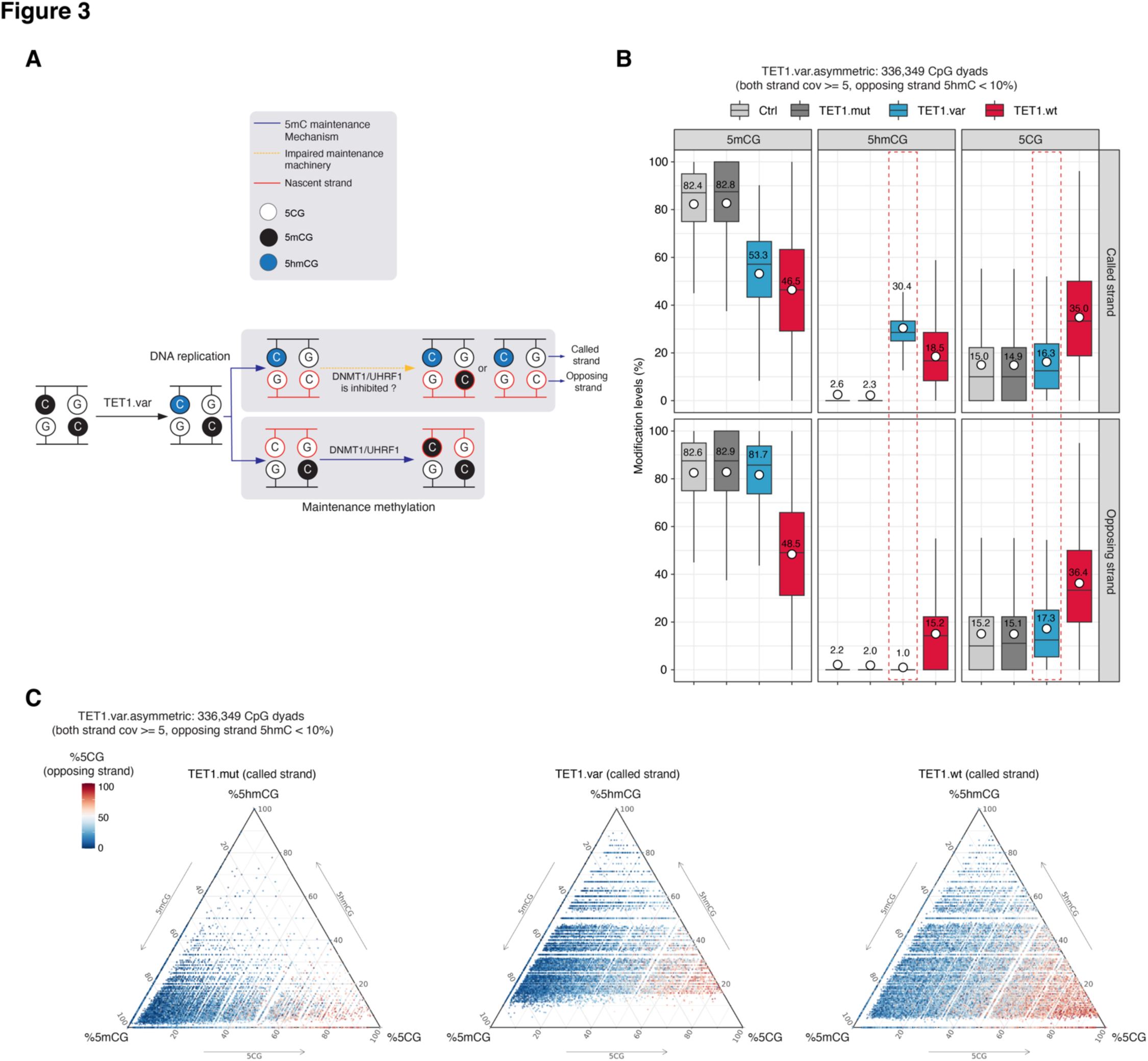
CpG sites enriched with 5hmCG resist 5mCG depletion on opposing strands in proliferating somatic. (**A**) Schematic diagram of the hemi-methylated CpG maintenance mechanism by DNMT1/UHRF1 and the potential impairment of hemi-hydroxymethylation on the CpG methylation maintenance on the opposing strand. **(B)** Box plots of 5mCG, 5hmCG and 5CG levels (%) on the called (top) and opposing (bottom) strands of asymmetrically hydroxymethylated CpG dyads (n = 336,349 sites; the population averages of 5hmCG and 5CG for this group of CpGs in TET1.var are highlighted by red dotted line) in TET1.var expressing cells. TET.mut/wt are shown as controls. **(C)** Ternary plots showing levels of 5CG, 5mCG and 5hmCG (%) on the called strand of asymmetrically hydroxymethylated CpG dyads (n = 336,349 sites) in TET1.var expressing cells. TET.mut/wt are shown as controls. The 5CG levels (%) on the opposing strand for the same CpG dyad are color coded (red: high; blue: low).

Because TET enzymes are known to exhibit distinct catalytic processivity at different *cis*-regulatory elements (Schutsky et al., 2018; Wu et al., 2014; Wu and Zhang, 2017), we next sought to determine whether 5hmCG generation by TET1.var can mediate active DNA demethylation at specific genomic features characterized by different levels of transcriptional activity or local chromatin states. To this end, we annotated major genomic features (e.g. promoters, enhancers and genic regions) with ENCODE histone modification profiles generated in wild-type HEK293T cells (Consortium, 2012). Across all annotated genomic features, TET1.wt but not TET1.var, induced a marked increase in 5CG levels (**Figure S3E**). Next, we performed quantitative analysis on a subset of genomic regions statistically enriched with comparable levels of 5hmCG in TET1.var and TET1.wt (middle panel in **Figure 2D**). As expected, transcriptionally active promoters and gene bodies are associated with lower levels of true 5mCG compared to their inactive counterparts; and both TET1.wt and TET1.var-expressing cells are associated with varying degrees of 5mCG oxidation across different genomic features (top panel in **Figure 2D**). When compared to TET1.mut, this analysis shows that TET1.wt induces marked increase in 5CG levels (mean 5CG restoration: 17.6%, ranging from 12.6% for active promoters to 22.1% for enhancers) at various regulatory elements. By comparison, TET1.var results in an approximately 10-fold less 5CG increase at corresponding genomic elements (mean 5CG restoration: 1.72%, ranging from 1.1% for active promoters to 2.6% for enhancers) (bottom panel in **Figure 2D**). Heatmap visualization of these 5hmCG-enriched genomic features further confirm that 5hmCG generation by TET1.var only leads to minimal changes in 5CG levels at these sites and their flanking regions (**Figure 2E**). Single-base resolution maps of 5mCG, 5hmCG and 5CG at representative genomic features also support the conclusion that 5mCG oxidation to 5hmCG by TET1.var is not sufficient to induce substantial 5CG restoration, suggesting that 5fCG/5caCG generation by TET1.wt and subsequent excision repair by TDG/BER are likely required for robust active DNA demethylation (**Figure 2F**).

### Hemi-hydroxymethylation is predominantly resistant to 5CG restoration on the opposing strand

Previous *in vitro* biochemical studies suggested that hemi-hydroxymethylation can substantially reduce the activity of maintenance methylation machinery (DNMT1/UHRF1) on the nascent strand of CpG dyads (**Figure 3A**), thus contributing to DNA demethylation in proliferative somatic cells. This model predicts that 5hmCG generation by TET1.var would result in a substantial increase in 5CG levels on the opposing strand in our system. To test this, we first performed strand-specific statistical calling of 5hmC-modified CG sites. Using a *P* value cut-off of 2.5 x 10^−4^, we identified a 4,183,293 and 928,205 5hmCG sites in TET1.wt– and TET1.var-expressing cells (**Figure S4A**). As anticipated, a substantially lower number of high-confidence 5hmCG sites were identified in TET1.mut controls (n=99,327). By restricting our analysis to a subset of 5hmC-modified CpG dyads with sufficient sequencing coverage on both strands (cov >=5), we observe that the majority of 5hmCG dyads in TET1.var (95.4%) are asymmetrically modified, whereas a higher number of symmetrically modified 5hmCpG dyads were detected in TET1.wt (TET1.var: 4.6% vs. TET1.wt: 18.2%), reflecting the higher catalytic activity of TET1.wt (**Figure S4B**).

We next analyzed 5mCG, 5hmCG and 5CG levels on both called and opposing strands for a cohort of asymmetrically modified CpG dyads catalyzed by TET1.var (mean 5hmC level=30.4% on called strand, blue in **Figure 3B**). Compared to TET1.mut control, this analysis shows that TET1.var only engenders a relatively small increase in 5CG levels on the opposing strand (mean: 2.2%; measured by the difference between TET1.var and TET1.mut), whereas TET1.wt induces a ∼10-fold higher increase in 5CG levels (mean: 21.3%; **Figure 3B**). Given the relatively low, but detectable, expression level of TET1-3 (**Figure S2C**), the endogenous TET activity may also contribute to detected 5CG restoration observed in TET1.var-expressing cells. Thus, this helps to quantitatively define the upper bound of TET1.var-mediated indirect depletion of 5CG on the opposing strand (up to 2.2%). At these hemi-hydroxymethylated CpG dyads, a marked decrease in 5mCG levels was only observed on the called strand (mean 5mCG oxidation: 29.5%; measured by the difference between TET1.var and TET1.mut), but not on the opposing strand (mean 5mCG oxidation: 1.2%) (**Figure 3B**), which suggests that 5hmCG (on the called strand) may largely be paired with 5mCG (on the opposing strand).

Similar analysis of a cohort of asymmetrically modified CpG dyads catalyzed by TET1.wt (mean 5hmC level=32.1% on the called strand, red in **Figure S4C)** indicates that 5CG levels on both called and opposing strands increase substantially in TET1.wt-expressing cells compared to TET1.mut. Thus, these results suggest that unlike the limited impact of steady-state hemi-hydroxymethylation (generated by TET1.var) on 5CG restoration on the opposing strand (**Figure 3A-B**), TET1.wt-mediated 5fC/5caC excision repair may induce the 5CG restoration on the called strand first, which subsequently causes replication-dependent 5mC loss on the opposing strand upon cell division (**Figure S4C-D**).

To extend beyond the analysis of population averages among asymmetrically modified CpG dyads, we directly linked all three CG modification states (5mCG/5hmCG/5CG in ternary plots) on called strands with levels of 5CG on opposing strands (colored coded: red – high 5CG levels, blue – low 5CG levels) in TET1.wt/mut/var-expressing cells (**Figure 3C** and **S4E**). The ternary plots reveal high levels of epigenetic heterogeneity among asymmetrically modified CpG dyads and show both 5CG and 5mCG preferentially exist in symmetrically modified configurations (5mC-modified on both strands in the left bottom corners, and unmodified 5CG on both strands in the right bottom corners in **Figure 3C** and **S4E**). Thus, these results suggest that asymmetrical 5hmCG on the called strand tends to be paired with 5mCG, but not 5CG, on the opposing strand. Taken together, our integrated base-resolution epigenomic analysis of all major CG modification states support a model in which hemi-hydroxymethylation of CpGs by TET1 stalling variant generally does not engender an indirect mode for 5CG restoration on the nascent strand upon DNA replication.

### Transcriptome-wide RNA expression analysis reveals gene regulatory roles for 5hmC as a *bona fide* epigenetic modification

A previous study demonstrated that a mouse Tet2 stalling variant is functionally distinct from wild-type Tet2 in the somatic reprogramming of mouse fibroblasts to pluripotent stem cells (Caldwell et al., 2021), suggesting these Tet2 catalytic processivity variants may regulate gene expression differently. However, as somatic reprogramming is a highly dynamic process for cellular state transitions, it becomes challenging to mechanistically dissect the specific gene regulatory roles of 5hmC generation and related DNA demethylation pathways in this paradigm that contains heterogenous cell populations. Thus, the ability to biochemically decouple 5hmC generation from 5fC/5caC excision repair and subsequent 5C restoration in proliferative HEK293T cells affords an opportunity for investigating the potentially divergent regulatory impact of wild-type and stalling mutant TET variants on gene expression in a more controlled cellular context.

Because exogenous 5hmC generated by stalling variant TET1.var in proliferative HEK293T cells is largely independent of both direct and indirect DNA demethylation in this relatively homogeneous cell population, we can specifically evaluate whether 5hmC can contribute to gene regulation as a *bona fide* epigenetic mark by performing a pairwise comparison between TET1.var and TET1.mut. Next, we reasoned that a pairwise comparison between TET1.wt and TET1.var can reveal the specific gene regulatory role of 5fC/5caC excision repair and/or unmodified cytosine restoration. Finally, comparing TET1.mut and control cells may uncover potential roles of catalytic activity independent functions of TET1 CD as a protein interacting scaffold. To this end, we performed total RNA sequencing (RNA-seq) to quantify expression levels of both protein-coding mRNAs and non-coding RNAs (ncRNAs) without polyA tails in control and TET1.wt/mut/var-expressing cells (**Figure S2A** and **Table S3**).

While the transcriptome of TET1.mut cells is largely similar to control cells (17 genes up, 130 genes down), both TET1.wt and TET1.var induced substantially more changes in gene expression compared to TET1.mut, displaying a roughly five-to-seven-fold larger set of differentially expressed genes (**Figure 4A**). Interestingly, protein-coding mRNAs reproducibly show both up– and down-regulation in response to TET1.var (5hmC generation alone) or TET1.wt (5hmC generation and 5fC/5caC excision repair) mediated epigenome-wide 5mC oxidation, whereas non-coding RNAs (annotated by Gencode database) tend to be down-regulated in TET1.var and TET1.wt cells (**Figure S5A-B**). Genes up-regulated by active DNA demethylation (up only in TET1.wt; *TPM2* in **Figure 4B**) and 5hmC generation alone (up in both TET1.wt and TET1.var; *ZCCHC12* in **Figure 4B**) are enriched for genes involved in pathways of specific cellular processes or tissue development and morphogenesis (left in **Figure S5C**). In contrast, down-regulated genes tend to be associated with molecular processes such as RNA splicing that require many functioning ncRNAs (*RUN4-1* in **Figure 4B**; right in **Figure S5C**).

**Figure 4.**
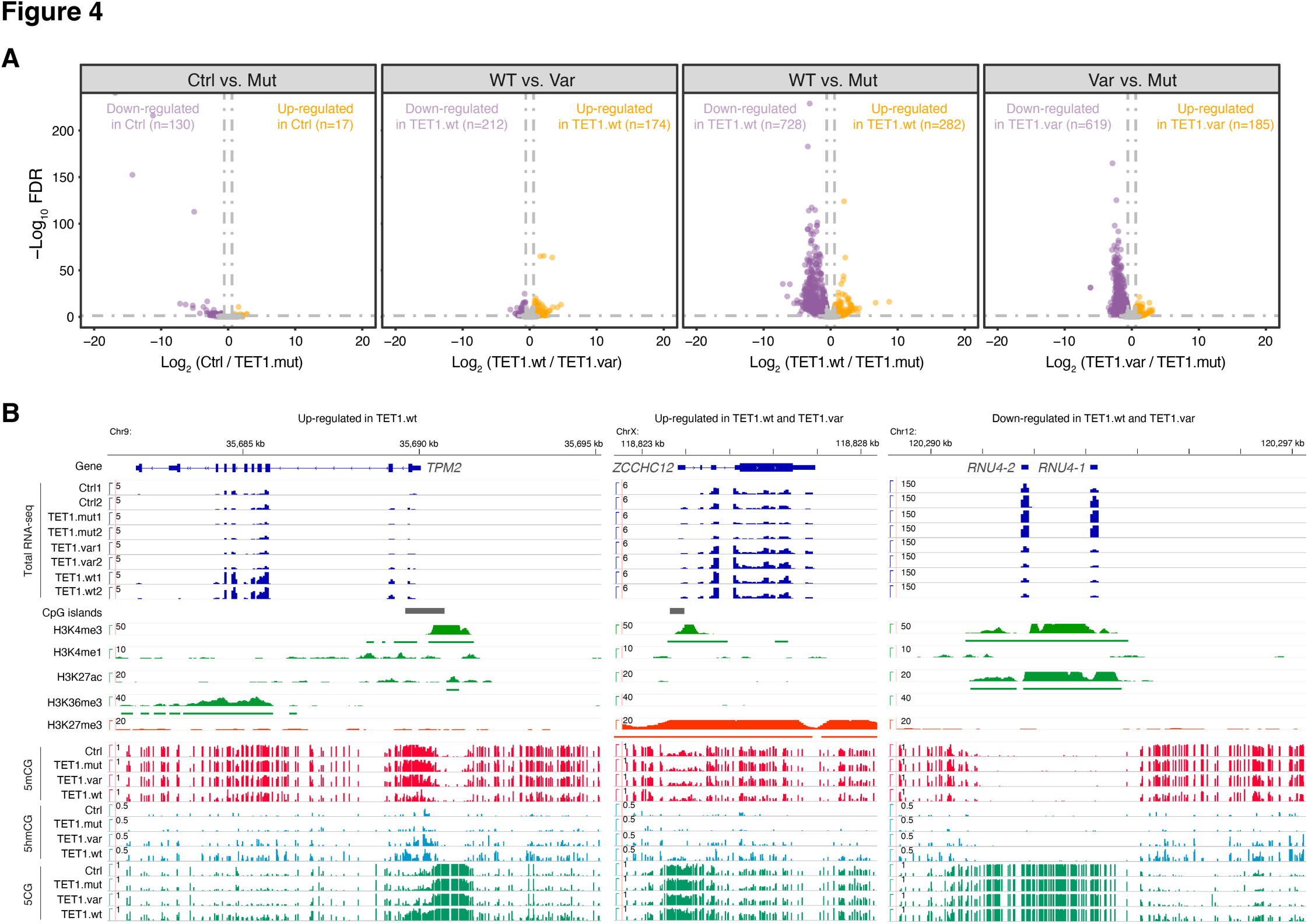
Total RNA sequencing analysis reveals 5hmC alone may act as a gene regulatory epigenetic cation. (**A**) Volcano plots of differentially expressed genes (DEGs, including both protein-coding and non-coding genes) among four pairwise comparisons: control versus TET1.mut (regulated by TET1 protein scaffold), TET1.wt versus TET1.var (regulated by iterative 5mC oxidation alone), TET1.wt versus TET1.mut (regulated by iterative 5mC oxidation and/or 5hmC generation), and TET1.var versus TET1.mut (regulated by 5hmC generation alone). The cutoff for identifying DEGs is FDR < 0.05 and fold change > 1.5. Up-regulated genes are shown in orange; down-regulated genes are displayed in purple; insignificant genes are shown in grey. **(B)** Genome browser tracks of gene annotations, CpG islands, major histone modifications (wild-type only, normalized signals to library size, counts per million reads), total RNA levels (CPM, counts per million reads; with both biological replicates shown), 5mCG (0-100%), 5hmCG (0-50%) and 5CG (0-100%) signals for three representative DEGs (up-regulated in TET1.wt: *TPM2*, up-regulated in TET1.wt/var: *ZCCHC12*, and down-regulated in TET1.wt/var: *RNU4*) in control, TET1.mut, TET1.var, and TET1.wt cells, with scale of y-axis shown.

## DISCUSSION

Since the initial discovery of TET-mediated 5mC oxidation to 5hmC and its unusually high enrichment in post-mitotic neurons (Kriaucionis and Heintz, 2009; Tahiliani et al., 2009), it has been speculated that 5hmC can potentially act as a *bona fide* epigenetic mark to regulate gene expression. Indeed, *in vitro* screening and biochemical characterization demonstrated that 5hmC can potentially either attract candidate reader proteins (Iurlaro et al., 2013; Spruijt et al., 2013) or repel 5mCG binding proteins such as *MECP2* (Kinde et al., 2015). However, TET enzymes initiate active DNA demethylation through either iterative 5mC oxidation and TDG/BER-induced excision repair, or by facilitating replication-dependent erasure of 5mC on the opposing strand by oxidized 5mC-mediated inhibition of maintenance machinery (DNMT1/UHRF1) activity or recruitment. Consequentially, it is technically challenging to bifurcate the direct (as a *bona fide* epigenetic mark) versus indirect (as an intermediate of TET-dependent active DNA demethylation pathways) contributions of 5hmC on gene regulation.

Here we sought to address two related questions regarding the epigenomic and gene regulatory functions of 5hmC by leveraging a human TET1 stalling variant to decouple the stepwise process of initial 5mC oxidation to 5hmC from subsequent 5fC/5caC generation/excision and repair. First, through developing a dCas9-SunTag based approach for targeted TET1.var recruitment and applying it to study an endogenous methylation-sensitive gene promoter, *RHOXF2B*, we demonstrate that TET1.var can robustly catalyze the initial oxidation of 5mC to 5hmC, but stalls at this step without further generating 5fC/5caC by using integrated epigenetic sequencing methods to disambiguate 5mC from 5hmC, and C from 5fC/5caC. Importantly, this result confirmed *in vivo* specificity of TET1.var catalytic processivity on an endogenous direct target. Second, we performed epigenome-wide editing using non-targeted TET1.wt/mut/var and harnessed paired (hydroxy)methylome sequencing to reveal that hemi-hydroxymethylated CpG dyads are not strongly linked to the restoration of 5CG across various genomic elements and on opposing strands in proliferative HEK293T cells. In contrast to the prevailing view, these quantitative epigenomic sequencing results provide cogent evidence that supports a model in which the direct mode of TET-dependent active DNA demethylation (through 5fC/5caC excision repair) is the predominant mechanism to restore 5CG, whereas the indirect pathway (through 5hmC-dependent inhibition of DNMT1/UHRF1) infrequently occurs in proliferative somatic cells. Finally, the total RNA-seq analysis of transcriptomic responses to epigenome-wide editing by TET1.wt and TET1.var reveals that 5hmC generated by TET1.var may induce similar levels of gene expression changes as those stimulated by TET1.wt. This suggests that in specific cellular contexts, 5hmC may act as a gene regulatory epigenetic modification without inducing substantial remodeling of the 5CG landscape at *cis*-regulatory regions.

In summary, our results suggest that 5hmC can not only act as an intermediate for active DNA demethylation involving 5fC/5caC excision repair, but also regulate gene expression as a relatively stable epigenetic modification in proliferative somatic cells. Moreover, the CRISPR/dCas9-based targeted epigenome-editing approach developed in this study provides a versatile toolkit for mechanistically dissecting of causal function of 5hmC versus 5fC/5caC excision repair at diverse regulatory elements in future studies.

### Limitations of study

HEK293T cells are highly proliferative but not synchronized in cell cycling in our study, so each cell may have gone through different number of cell cycles, which may complicate the quantitative interpretation of the population average of our epigenomic sequencing results in terms of estimating the theoretical upper bound of 5hmC-induced passive DNA demethylation. Furthermore, because these cells also express *de novo* DNA methyltransferases, DNMT3A and DNMT3B, at low levels, we cannot exclude the possibility that the lack 5CG restoration in TET1.var-expressing cells is in part due to activity of DNMT3A/3B after cell division. Finally, TET CDs may also potentially modulate RNA abundance by directly modifying RNAs, so the observed TET1 catalytic activity-dependent changes in gene expression may in part be regulated at the RNA level, instead of the DNA level.

## STAR METHODS RESOURCE AVAILIBILITY

### Lead Contact

Further information and requests for resources and reagents should be directed to and will be fulfilled by the Lead Contact, Hao Wu.

### Materials Availability

Plasmids generated in this study will be deposited to Addgene.

### Data and code availability

All sequencing data associated with this study will be available on the NCBI Gene Expression Omnibus (GEO) database upon publication. The analysis source code underlying the final version of the paper will be available on GitHub repository (https://github.com/wulabupenn/) upon publication.

## EXPERIMENTAL MODEL AND SUBJECT DETAILS

### Human HEK293T cell culture

Human HEK293T cells were maintained at 37°C with 5% CO_2_ in CO_2_ incubators (Thermo Scientific, 51-033-774) in Dulbecco’s Modified Eagle’s Medium (DMEM) (Gibco, 11965084), supplemented with 10% fetal bovine serum (Gibco, 16000044), 5% L-glutamine (Gibco, 25030081), 5% nonessential amino acid (Gibco, 11140050), and 5% sodium pyruvate (Gibco, 11360070) in 10-cm dishes and passaged every 2-3 days at 70% confluency with PBS and 0.05% Trypsin-EDTA (Gibco, 25300054).

## METHOD DETAILS

### Generation of Suntag-TET1.wt/var/mut constructs, and all-in-one plasmids

Catalytically inactivated hTET1CD (H274Y/D276A) gBlock was ordered from Integrated-DNA technologies (IDT) and amplified using 2× KAPA HiFi PCR kits, Hotstart Ready mix (Kapa biosciences, KK2602) per the manufacturer’s instruction. Site directed mutations to generate the 5hmC-stalling Suntag-TET.var (T246E) was introduced by amplifying the previously validated Suntag-TET1CD construct (Addgene #82559) with mutation containing primers (IDT). 1ug Suntag-TET1CD was digested with Nhe1-HF (New England Biolabs, R3131L) and Not1-HF (New England Biolabs, R3189S) to isolate the backbone and release TET1CD. Digested backbone and TET1.mut/var amplified fragments were size-selected with Qiaquick Gel Extraction kit (Qiagen, 28506), and cloned into Suntag-TET1CD Nhe1-HF/Not1-HF digested backbones with NEBuilder HiFi DNA assembly Master Mix (New England Biolabs Inc. E2621L). Following bacterial transformation into Stbl3 competent cells, inoculation, and construct isolation with the NucleoSpin Plasmid kit (Macherey-Nagel, REF 740588.250), plasmids were verified with sanger sequencing.

To generate All-in-one plasmids that express *RHOXF2B* sgRNA concomitantly, 1ug SunTag-(TET1.wt/mut/var) constructs were linearized with Af1II (New England Biolabs, R0520S), and isolated with the Zymo DNA clean & concentrator kit (Zymo Research, D4033). sg*RHOXF2B* gblocks were ordered from IDT, and cloned into digested backbones and isolated as described above. Plasmids were subsequently validated with sanger sequencing.

### Construction of sgRNA expressing vectors with Golden Gate Cloning

Complimentary *RHOXF2B* sgRNA oligos were ordered from IDT. 100uM Oligos were annealed using 10× T4 DNA ligase reaction buffer (New England Biolabs, B0202S), and T4 Polynucleotide Kinase (New England Biolabs, M0201L). 1uL of 1uM annealed sgRNA oligos were then cloned into 30ng Lenti-sgRNA(MS2)-puro constructs (Addgene #73797) through Golden-gate assembly with BsmBI (Thermo Scientific, FD0454), 2× rapid ligase buffer (Enzymatics, B101), Bovine Serum Albumin (New England Biolabs, B9000S), and T7 ligase (Enzymatics, L602L). Constructs were transformed, inoculated, isolated, and sequenced as described above.

### Transient transfection and cellular expansion

For FACS-sorted All-in-one experiments, 2ug of Suntag platform containing scrambled or *RHOXF2B* sgRNA expressing constructs, and pTY-GFP (transfection control) were transiently transfected in suspension with dissociated 2×10^6^cells per sample in Opti-MEM I Medium (Thermo Fisher Scientific, 31985062), with Lipofectamine 2000 (Invitrogen, 11668019) per manufacturer’s recommendation into 12-well plates. However, Lipofectamine 2000-DNA complexes were incubated for 45 minutes as opposed to manufacturer’s instruction. 24-hours post-transfection, the cells were expanded onto-6 well plates and incubated for an additional 48-hours prior to GFP+/DAPI-gated FACS followed by gDNA/RNA (Zymo Research, D7001), and protein isolation.

For unsorted experiments, 1×10^6^ dissociated cells were co-transfected in suspension with 800ng of Suntag/pTY-GFP, and 200ng of scrambled or *RHOXF2B* sgRNA expressing plasmids from the generated Lenti-sgRNA(MS2)-puro constructs. After 24 hours, cells were expanded, and further incubated an additional 24hours before 0.5 µg/ml puromycin was administered to drug-select for sgRNA expressing transfected cells. After 24 hours, gDNA and RNA was isolated with Quick-DNA/RNA miniprep kits (Zymo Research, D7001).

### Flow Cytometry to isolate for GFP+/DAPI-transfected cells

Transfected cells are washed with PBS and dissociated with 0.05% Trypsin-EDTA, prior to being resuspended with media, and spun down to form a pellet. Cells are then resuspended with 0.5mL 0.1% PBS-BSA solution and passed through a 40uM filter (Fisher Scientific, 08-771-1) to dissociate into single-cell suspension. The micron filter is subsequently washed with another 0.5mL 0.1% PBS-BSA solution. 1uL 100mg/uL DAPI (Sigma Aldrich, D9542) was added to each sample to stain for compromised cellular membranes. Cell suspension was then sorted with BD Biosciences influx cell sorter at the University of Pennsylvania Flow Cytometry and Cell Sorting core for DAPI negative, and GFP positive cell populations into a 15mL conical tube loaded with 0.5mL PBS.

### Hydroxymethylated DNA immunoprecipitation (5hmC-DIP) qPCR

250 ng gDNA was sheared into 250 bp fragments using a M220 Focused ultra-sonicator (Covaris) with 50 peak power, 20 duty power, and 200 cycles, for 120s. Dynabeads Protein G (Thermo Fisher Scientific, 10004D) were washed with 1mL 0.5% BSA PBS blocking buffer twice on a magnetic rack and then resuspended with 0.5% BSA PBS blocking buffer at the original volume. 1uL (1mg/ml) of anti-5hmC antibody (Active Motif, 39769) or IgG was added for each reaction, and rotated for 5 hours in 4°C. Upon completion, fragmented gDNA is combined with 5x MeDIP IP buffer (14% 5M NaCl, 5% 1M Sodium phosphate Buffer, 2.5% 10% 10xTriton) in a separate PCR strip. 10uL of this mix is removed for benchmarking input. The antibody-Dynabead mix is then added at a 1:1 ratio to the gDNA-MeDIP IP buffer mix in the PCR strip, and undergoes overnight immunoprecipitation at 4°C. Samples were washed with 200ul 1x MeDIP IP buffer, three times on a magnetic rack over ice. 62.5uL MeDIP digestion buffer, and 0.875uL (20ug/uL) Proteinase K is added to each sample, and incubated in the thermal cycler at 55°C for one hour. Eluate is transferred to a new PCR strip, where it is purified with RNA Clean XP beads (Beckman Coulter Life Sciences, NC0068576) at 2x volume with standard instructions. qPCR analysis was subsequently performed with 400nM primers (IDT), and 2x PowerUp SYBER Green master mix (Thermo Fisher Scientific, A25742), to evaluate pulled-down methylated DNA fragments enrichment.

### Western blot

FACS-sorted GFP+/DAPI-HEK293T cell populations were first washed with cold PBS, and lysed with 10X RIPA buffer (Cell Signaling Technology, 9806) supplemented with 50x Complete protease inhibitor cocktail (Sigma Aldrich,11873580001). Lysates were centrifuged in 4°C, and the supernatant was transferred to a new tube on ice. Protein concentration was measured using Bio-rad DC protein assay (Bio-rad, 5000112) and a Beckman Spectrometer. 5ug of protein per sample was denatured with 4x Laemmli buffer (Bio-rad, 1610747) for 10 minutes at 95°C, and ran on a 4-15% Mini-PROTEAN-TGX precast protein gel (Bio-rad, 4561085) at 165V. For TDG overexpression experiments, 20 ug protein was used as starting material. The protein is then transferred to PVDF midi membranes using the Trans-blot Turbo Transfer System (Bio-rad, 1704150).

Upon transfer completion, the PVDF membrane is washed with 1x TBST buffer (Thermo Scientific, 28358), and then blocked with 5% milk 1xTBST solution at room temperature for 1 hour shaking. The membrane is washed again with 1x TBST solution, and then incubated the primary antibody solution (3% BSA in TBST, 1:250 anti-HA (Sigma-Aldrich, A2095-1ML), 1:500 anti-FLAG (Sigma-Aldrich, F1804-1MG), 1:3000 anti-ACTB (Cell Signaling Technology, 3700S) overnight in 4°C while shaking. PVDF membrane is then washed with 1x TBST solution, and blocked with a secondary antibody solution (1:500 antibody dilution) comprised of 5% milk and 1xTBST for 1 hour while shaking. The membrane is washed one final time with 1x TBST, and then incubated with Clarity Max ECL Western Blotting Substrates (Bio Rad, 1705060), prior to visualization with a Bio Rad ChemiDoc Imaging System.

### Gene expression analysis with RT-qPCR

500ng RNA isolated from cell populations are synthesized into cDNA using iScript Reverse Transcription Supermix (Bio Rad, 1708841) per manufacturer’s recommendations. The resulting solution is further diluted 1:2.5 with ddH_2_O for downstream qPCR. Converted cDNA is quantified using 2x KAPA SYBR Fast Master Mix ROX Low (KAPA Biosystems, KK4620), with primers against upstream *RHOXF2B* TSS, and *ACTB* as a baseline, to evaluate relative changes in gene transcription.

### Bisulfite conversion and A3A deamination for locus-specific BS-seq and bACE-seq

To generate the requisite phage spike-in controls to evaluate A3A deamination efficiency, lambda phage DNA was *in vitr*o CpG methylated using the M.SssI methyltransferase (New England Biolabs, M0226M) with S-adenosylmethionine (SAM) for 2 hours in 37°C. After the initial incubation, additional enzyme and SAM is added to the solution for another 4 hours, prior to being inactivated at 65°C for 20 minutes. The phage DNA is then purified using 1.6x volume of homebrewed solid phase reversible immobilization (SPRI beads) (1mL Sera-Mag SpeedBeads (GE Healthcare, GE1715210401150), 9g PEG 8000 (Sigma, 1546606), 10mL 5M NaCl, 500µL 1M Tris-HCl pH 8.0 (Thermo Fisher Scientific, 15568025), 100µL 0.5M EDTA pH 8.0 (Thermo Fisher Scientific, R1021). M.SssI *in vitro* methylated lambda phage spike-ins are added to each sample at 1% total mass gDNA converted.

10 ng of gDNA isolated from GFP+/DAPI-FACS-sorted cell populations was first spiked with lambda phage controls and was bisulfite-converted with the Zymo EZ DNA Methylation Direct Kit (Zymo Research, D5020) per the manufacturer’s protocol, with exception of using an alternate elution buffer. In lieu of the manual’s recommendation, 15uL 1mM Tris-HCl pH 7.5 elution buffer is used to maintain A3A reaction conditions around pH 6.0. For 5uL of eluate, 1uL of 20mM 10x MES pH 6.0 and 0.1% tween, and 1uL DMSO is added to solution over ice. Samples are then denatured in a thermal cycler at 95°C for 5 minutes, and immediately transferred to a pre-chilled PCR rack in a –80°C fridge to snap-cool. 1uL of (80uM) A3A purified as previously described, 1uL of 20mM 10x MES pH 6.0, and 2uL of ddH_2_O is added to each sample prior to thawing, and incubated at 37°C for 2 hours. Converted samples are then bead purified with 1.6x volume homebrew SPRI beads, and eluted with 20uL of 10mM Tris-HCl pH 8.5.

### MAB-seq in vitro methylation and bisulfite conversion

0.5% unmethylated lambda phage spike-ins are added to 100 ng of gDNA isolated from GFP+/DAPI-FACS-sorted cells to serve as internal controls for *in vitro* CpG methylation efficiency in each sample. During the first round of *in vitro* methylation, 10x Mg^2+^-free buffer, 0.64 mM SAM, and 0.8 units/uL of M.SssI methyltransferase is added to each sample and incubated for 4 hours 37°C. After completion, an additional 0.64 mM SAM and 6 units of M.SssI enzyme is added to the reaction, and incubated for 8 hours at 37°C, and then inactivated with 20 min incubation at 65°C. DNA is then isolated using phenol-chloroform:isoamyl alcohol (25:24:1) (Thermo Fisher Scientific, 15593031) as previously described (Wu et al., 2016). The gDNA is eluted with 25.2uL ddH_2_O and undergoes a second round of *in vitro* CpG methylation with previously described parameters above with exception that 10x Mg^2+^-free buffer is replaced with equal volume of NEB Buffer #2 (New England Biolabs, B7002S). The reaction is incubated again for 4 hours at 37°C, and another 8 hours after addition of 0.64 mM SAM and 6 units of M.SssI enzyme. M.SssI is then deactivated with 20 minutes of incubation in 65°C. gDNA is isolated with phenol-chloroform extraction as previously described, and eluted with 20uL of ddH_2_O. Samples were bisulfite converted using the Qiagen Epitect DNA Bisulfite Kit (Qiagen, 59104) per manufacturer’s instructions. However, thermal cycling conditions were altered so that the parameters were repeated twice for a total of 10 hours. Samples were purified with the manual’s instructions, and eluted in 30uL of 10mM Tris-HCl pH 8.5.

### Locus-specific library preparation and sequencing

All reactions were performed in a sterile, ventilated hood to limit contamination from the environment. BS/bACE/MAB-seq converted gDNA is amplified with 500nM of primers against *RHOXF2B* TSS containing adapters for downstream sample indexing, with KAPA2G Robust kits (KAPA Biosystems, KR0379). An additional set of primers against spike-in lambda phage DNA is performed concomitantly to evaluate the effects of A3A deamination (bACE) against 5mC, or *in vitro* CpG methylation (MAB) on unmodified C in each sample. Thermal cycling conditions are as recommended from the manufacturer. However, individual samples were optimized by modification to gDNA amplified, primer concentration, cycles of amplification, and reaction volume to ensure robust locus amplification devoid of environmental contamination. PCR Adapted samples are then gel purified using the Qiquick Gel Extraction kit. 1ng of each sample is then uniquely indexed using the NEBNext Multiplex Oligos (New England Biolabs, E7335L, E7500L) with 2x KAPA HiFi Hotstart Ready mix for 8 cycles of amplification with the protocol’s instructions. The indexed fragments are then isolated using the Qiquick Gel Extraction kit. The 4nM library is pooled together, and its size is evaluated with the Bioanalyzer (Agilent, 5067-4626), and pair end sequenced on the Illumina Miseq using the MiSeq Reagent Nano Kit v2 (300-cycles) (Illumina, MS-102-2002).

### Integrated whole genome BS-seq and bACE-seq

The whole genome BS-seq (WGBS) and bACE-seq (WG-bACE-seq) experiments were performed as previously described with minor modifications (Fabyanic et al., 2023; Fabyanic et al., 2021). Briefly, ∼15 ng of genomic DNA from each sample was first spiked in with *in vitro* methylated lambda phage genomic DNA (0.2%) as controls and was then subjected to bisulfite conversion (EZ DNA Methylation-Direct Kit, Zymo Research Cat# D5020). Half of the bisulfite converted DNA was used for low input WGBS analysis (for 5mC+5hmC profiling). The other half of eluate was subjected to the low input WG-bACE-seq workflow (for 5hmC profiling). For each bACE-seq reaction, 1.5µL 200mM MES pH 6.0 + 0.1% Tween and 1.5µL DMSO were added to the 9μL eluent. The samples were then denatured at 95°C for 1min and snap cooled by transfer to a PCR tube rack pre-incubated at −80°C (for bulk samples). Before thawing, 1.5µL 200 mM MES pH 6.0 + 0.1% Tween-20 and 1.5µL 5μM A3A were added to each reaction to a final volume of 15µL (for a final concentration of 500nM/µL A3A per reaction). The deamination reactions were incubated at 37°C for 2h, purified with 1.6x homebrew SPRI beads, eluted in 9µL Low EDTA TE buffer.

To add the first PCR adaptor (P5), random priming reactions were performed for both WGBS and WG-bACE-seq library preparation. Deaminated DNA was first heated at 95°C using a thermocycler for 3min to denature and were immediately chilled on ice for 2min. 10µL enzyme mix (2µL Blue Buffer (Enzymatics B0110), 1µL 10mM dNTP (NEB N0447L), 1µL Klenow exo (50U/µL, Enzymatics P7010-HC-L), and 6µL water) was added to each well and reactions were mixed by vortexing. Plates or reactions were treated with the following program using a thermocycler: 4°C for 5min, ramp up to 25°C at 0.1°C/sec, 25°C for 5min, ramp up to 37°C at 0.1°C/sec, 37°C for 60min, 4°C forever. Following this, 2μL Exonuclease 1 (20U/μL, Enzymatics X8010L) and 1μL Shrimp Alkaline Phosphatase (rSAP) (1U/μL, NEB M0371L) was added to each reaction followed by vortexing and incubation in a thermocycler at 37°C for 30min followed by 4°C forever.

To add the second PCR adaptor (P7), the reactions were denatured in a thermocycler at 95°C for 3 min and subsequently chilled on ice for 2 min. 10.5μL Adaptase master mix (2μL Buffer G1, 2μL Reagent G2, 1.25μL Reagent G3, 0.5μL Enzyme G4, 0.5μL Enzyme G5, and 4.25μL Low EDTA TE buffer; Accel-NGS Adaptase Module for Single Cell Methyl-Seq Library Preparation, Swift Biosciences 33096) was added to each reaction, followed by vortexing. Reactions were incubated in a thermocycler at 37°C for 30min then 4°C forever. Subsequently, 30μL PCR mix (25μL KAPA HiFi HotStart ReadyMix, KAPA BIOSYSTEMS KK2602, 1μL 30μM P5 indexing primer, and 5μL 10μM P7 indexing primer) were added to each well, followed by mixing with vortexing.

Next, we perform qPCR to determine the optimal cycle number of amplification for indexing PCR. Reactions were transferred to a thermocycler programmed with the following stages: 95°C for 2min, 98°C for 30sec, 12-15 cycles of [98°C for 15sec, 64°C for 30sec, 72°C for 2min] (optimal cycle number may vary between samples), 72°C for 5min, and 4°C forever. PCR products were cleaned with two rounds of 0.8x homebrew SPRI beads, concentration was determined via QbitⓇ dsDNA High Sensitivity Assay Kit (Invitrogen Q32851), and library size and quality was determined via Bioanalyzer (Agilent High Sensitivity DNA Kit, 5067-4626). Libraries were first sequenced on an Illumina MiSeq using the 300-cycle kit (v2) to determine the WGBS and WG-bACE-seq library quality. Final libraries were diluted and pooled together for sequencing on Illumina NovaSeq 6000 using a 300-cycle High Output v2 Kit (150bp x 2).

### Bulk RNA sequencing

RNA was extracted from GFP+/DAPI-FAC-sorted transient transfected HEK293T cells with ZR-Duet DNA/RNA MiniPrep Kit (Zymo Research, D7001). RNA-seq libraries were prepared using SMARTer Stranded total RNA-seq Kit v3 (Clontech, 634486) per the manufacturer’s instructions. However, 20ng of RNA was used as starting input compared to the recommended 10ng. Library concentration and complexity was validated with a Qubit Fluorometer and Agilent Bioanalyzer 2100 (Agilent, 5067-4626). Libraries were diluted to 4nM and pooled together for sequencing on Illumina NextSeq 500 using a 300-cycle High Output v2 Kit.

## QUANTIFICATION AND STATISTICAL ANALYSIS

### Read mapping and quality filtering whole-genome BS-seq and bACE-seq

The pre-processing (read alignment, quality filtering and read deduplication) was performed for BS-seq and bACE-seq datasets as previously described with minor modifications (Fabyanic et al., 2023; Fabyanic et al., 2021). Briefly, demultiplexing of inline barcodes was first performed allowing up to 1-nt mismatch. The data quality was examined with FastQC (http://www.bioinformatics.babraham.ac.uk/projects/fastqc/). Raw sequencing reads were trimmed for adaptor sequences and inline barcodes using Cutadapt (Kechin et al., 2017) with the following parameters in paired-end mode: –f fastq –q 20 –u 16 –U 16 –m 30 –a AGATCGGAAGAGCACACGTCTGAAC –A AGATCGGAAGAGCGTCGTGTAGGGA. The trimmed R1 and R2 reads were mapped independently against the reference genome (mm10) using Bismark (Krueger and Andrews, 2011) (v0.18.2) with following parameters: –-bowtie2 –D 15 –R 2 –L 20 –N 0 –score_min L,0,-0.2 (––pbat option was turned on for mapping R1 reads). Uniquely mapped reads were filtered for minimal mapping quality (MAPQ>=10) using samtools (Li et al., 2009). PCR duplicates were removed using the Picard *MarkDuplicates* (http://broadinstitute.github.io/picard/). To eliminate reads from strands not deaminated by A3A, reads with three or more consecutive non-converted cytosines in the CH context were removed using *filter_non_conversion* in Bismark. Base calling of unmethylated and methylated cytosines was performed by *bismark_methylation_extractor* in Bismark in each individual nucleus. 5hmC signals were calculated as % of C/(C+T) at each cytosine base. Sequencing reads for WG-bACE-seq were pre-processed as previously reported (Schutsky et al., 2018).

### Statistical calling of 5hmC-enriched genomic regions or CpGs in whole-genome bACE-seq datasets

For each genomc regions (**Figure 2B**) or CG dinucleotides (**Figure S4A**), we counted the number of ‘C’ bases from bACE-seq reads as 5hmC (denoted *N_C_*) and the number of ‘T’ bases as methylated or unmodified cytosines (denoted *N_T_*). For statistical calling, we used the binomial distribution (*N* as the sequencing coverage (*N_T_* + *N_C_*) and *p* as the error rate of A3A deamination (1.61%, averaged non-conversion rate for 5mCG in spiked-in λ phage DNA, from eight independent measurements)) to assess the probability of observing *N_C_* or greater by chance. We then merged the two replicates and performed statistical calling of 5hmCG enriched 10-kb genomic regions or CpG sites (*P* < 2.5 x 10^−4^) using a binomial distribution model previously established for identifying 5hmC-modified CpG sites in mammalian genomes (Schutsky et al., 2018). To filter out low quality regions or CpG sites for statistically calling, we restricted our statistical analysis to CG sites covered by at least 200 reads per region (*n* = 282,870 10-kb genomic intervals in **Figure 2B**) or 5 reads per strand (*n* = 21,392,782 CpG sites in **Figure S4A**).

### Calculating the true level of 5mCGs by combining bACE-seq with BS-seq

For each CG site, the levels of 5mC and 5hmC were estimated using the MLML tool (Qu et al., 2013). This approach arrives at maximum likelihood estimates for the 5mC and 5hmC levels by combining data from bACE-seq and BS-seq (see below). Only CG sites with 0 conflicts were considered for further analysis. From the MLML output, the level of unmodified CG was estimated by [100% − (abundance of 5hmC + 5mC)]. The results were further filtered, such that 5CG, 5mCG, and 5hmCG levels were non-negative. For generating ternary plots (**Figure 3C** and **Figure S4E**), levels of 5CG, 5mCG, and 5hmCG (as percentage of the sum of [CG + 5mCG + 5hmCG]) were calculated within CpG dyads across the genome.

### Total RNA sequencing analysis

Raw data (fastq files) were mapped with strand-specific and single-end mode by Hisat2 (v2.2.1) (Kim et al., 2019). Unique mapped reads were kept with the following parametters (–F 4 –F 256) and sorted by samtools (v1.7) (Li et al., 2009). Human annotation file (Gencode v39) was downloaded from gencode. featureCounts (v2.0.1) (Liao et al., 2014) was used to quantify reads from exon. Differential expression analysis of protein coding genes (n = 19,986) and non-protein coding genes (n = 41,547) was performed using the edgeR (v3.30.3) (Robinson et al., 2010) R package with the cutoff (FDR < 0.05 and | fold change | > 1.5). Heatmap of differentially expressed genes was visualized by ComplexHeatmap (v2.11.2) R package (Gu, 2022).

Gene ontology gene sets were retrieved from msigdbr (v7.5.1) (Dolgalev, 2020) R package. Hypergeometric test of significant pathways (P.adjust < 0.1) was performed using enricher function from clusterProfiler (v3.18.1) (Yu et al., 2012) R package.

### Data visualization

Plots were generated using the ggplot2 (v. 3.3.0), and packages in R (version 3.5.1). RT-qPCR bar plot was generated with GraphPad prism (v9.1.0). We used Integrative Genomics Viewer (IGV, v2.11.4) to visualize WG-BS-seq and WG-bACE-seq signals using hg38 Refseq transcript annotation as reference (**Fig. 2** and **Fig. 4**). 5hmCG signals (both strands combined) are indicated by upward ticks, with the height of each tick representing the fraction of modification at the site ranging from 0-50%.

### Statistics

Statistical analyses were performed using R. Statistical details for each experiment are also provided in the figure legends. No statistical methods were used to predetermine sample size for any experiments. All group results are expressed as mean +/− standard deviation unless otherwise stated. Specific p-values used for calling modified cytosine bases are explicitly stated in the text and figure legends. Each figure legend explicitly states the number of independent experiments.

### Published data sets

For **Figure 2D-F**, **S3E**, and **4B**, we used the following published data sets: H3K4me1 (GSE174861), H3K27ac (GSE174866), H3K4me3 (GSM945288), H3K36me3 (GSE175320), H3K27me3 (GSE133391).

For data initially mapped to hg19, they were re-mapped to hg38 using liftOver. Genomic coordinates for exon, intron and gene body of UCSC RefSeq genes (GRCh38) were downloaded from Table browser (https://genome.ucsc.edu).

## SUPPLEMENTAL INFORMATION

**Figure S1**. Validation experiments for the CRISPR/dCas9-SunTag based 5hmC editing system. Related to Figure 1.

**Figure S2**. Experimental design and expression analysis for genome-wide epigenome editing in HEK293T cells. Related to Figure 2.

**Figure S3**. Validation experiments for integrated whole genome BS-seq and bACE-seq analyses. Related to Figure 2.

**Figure S4**. TET-mediated iterative 5mC oxidation is required for active DNA demethylation on both called and opposing strands of CpG dyads in proliferating somatic cells. Related to Figure 3.

**Figure S5**. Total RNA sequencing analysis reveals the gene regulatory roles of TET-mediated stepwise, iterative 5mC oxidation on protein-coding and non-coding RNAs. Related to Figure 4.

**Table S1**. Oligonucleotides used in this study

**Table S2**. Whole genome BS-seq and bACE-seq libraries constructed in this study

**Table S3**. RNA-seq libraries constructed in this study

## ACKNOWLEDGEMENTS

We are grateful to all members of the Wu lab for helpful discussion. This work was supported by the Penn Epigenetics Institute, the National Human Genome Research Institute (NHGRI) grants R01-HG010646 and U01-HG012047 (to H.W.) and NHGRI grant F31HG011429 (to A.W.).

## AUTHOR CONTRIBUTIONS

Conceptualization: AW, HW

Methodology: AW, PH, QQ, EBF, HW

Investigation: AW, HW

Bioinformatic Analysis: AW, HZ, HW

Funding acquisition: AW, HW Writing: AW, HW

Supervision: HW

## DECLARATION OF INTERESTS

Authors declare that they have no competing interests.

## INCLUSION AND DIVERSITY

We support inclusion, diverse and equitable conduct of research.

---

**Figure S1.**
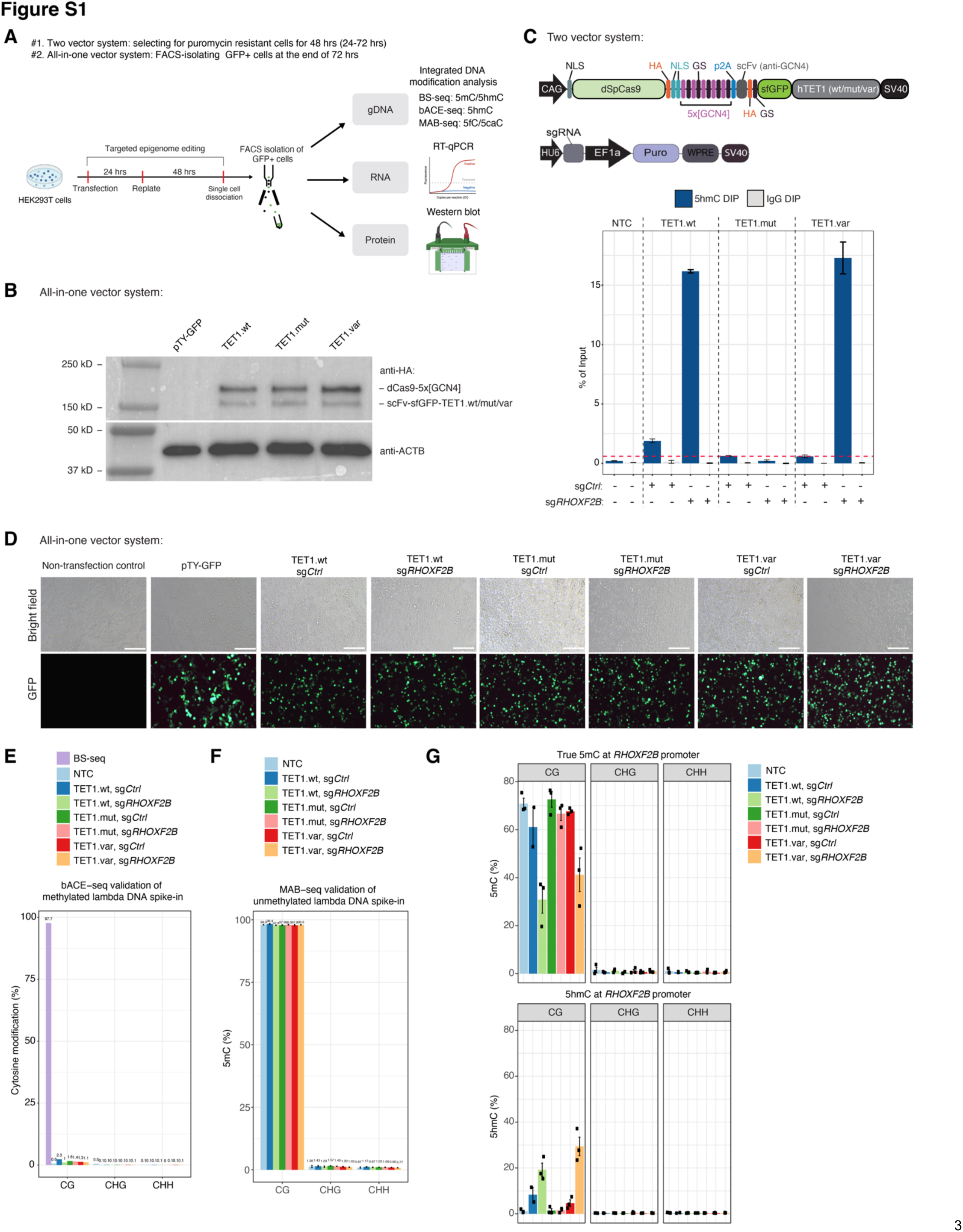
Validation experiments for the CRISPR/dCas9-SunTag based 5hmC editing system. (**A**) Schematic experimental workflow for transfecting SunTag.TET (wt/mut/var) constructs into HEK293T cells. All-in-one plasmids include the expression of the sgRNA in the same construct. Non-inclusive transfection experiments utilized co-transfection strategies outlined in the methods. Following transfection, cells are sorted with FACS to isolate GFP+/DAPI-cell populations. gDNA, RNA, and protein are isolated for downstream integrated DNA modification-, transcription-, and protein analysis. Abbreviations: FACS, Fluorescent Activated Cell Sorting; BS-seq, Bisulfite Sequencing; bACE-seq, bisulfite-assisted APOBEC Coupled Epigenetic Sequencing; MAB-seq, M.SssI Assisted Bisulfite Sequencing; scFv, single-chain variable Fragment; sfGFP; Super Folder Green fluorescent Protein; ACTB, β-Actin. **(B)** Representative western blot analysis evaluating the protein expression of our recombinant plasmids in FACS-sorted GFP+/DAPI-cells. Top bands represent dCas9-5x[GCN4]; bottom bands represent scFV-sfGFP-TET1.wt/mut/var. β-Actin is used as a loading control to ensure commensurate sample loading across samples. **(C)** (Top) Schematics of the two-vector system, where dCas9-SunTag-TET1 and sgRNA are encoded by separate vectors. (Bottom) 5hmC DNA immunoprecipitation (5hmC-DIP) in puromycin-selected HEK293T cells qualitatively evaluating SunTag.TET1.var 5hmC generation capacity relative to SunTag.TET1.wt and SunTag.TET1.mut at the *RHOXF2B* promoter. Enrichment is measured by qPCR as a percentage relative to input. Red dotted lines denote average background levels of 5hmC from sg*Ctrl* samples. **(D)** Brightfield (top) and fluorescent (bottom) microscopic visualization of transfection efficiency measured by GFP signals. Abbreviations: sg*Ctrl*, scrambled sgRNA; sg*RHOXF2B*, *RHOXF2B* promoter targeting sgRNA. pTY-GFP, a lentiviral vector encoding the EF1a promoter and EGFP transgene. **(E)** bACE-seq samples are individually spiked with M.SssI *in vitro* CpG methylated lambda phage DNA to benchmark APOBEC3A (A3A) 5mC deamination efficiency. A3A-mediated deamination is measured for CG/CHG/CHH sequence contexts. CHG/CHH 5mC is further utilized to evaluate for off-target *in vitro* methylation from M.SssI. BS-seq samples (furthest left in purple) do not experience A3A enzymatic deamination to benchmark M.SssI *in vitro* methylation efficiency against unmethylated lambda phage DNA. See methods for details on generating spike-in controls. Numbers on top represent mean percentage of called cytosine modifications. Percentages are calculated as total cytosine modification calls/total CpG sites across the amplicon multiplied by 100. **(F)** MAB-seq samples are individually spiked in with unmethylated lambda phage DNA to benchmark M.SssI *in vitro* methylation against unmodified CpG dyads to protect against BS-mediated deamination. Unmodified C protection is quantified by measuring total called C/C+T multiplied by 100 as a percentage in the CpG context. CHG/CHH sequences are measured to quantify off-target M.SssI *in vitro* methylation effects. **(G)** Locus-wide quantifications of true 5mC (top) and 5hmC levels (bottom) in CG and non-CG (CHG/CHH) sequence contexts. True 5mC levels are calculated by subtracting called cytosine modifications from bACE-seq from BS-seq quantities. Bar heights represent mean values. Individual dots represent independent biological replicates. Error bars denote +/− standard error.

**Figure S2.**
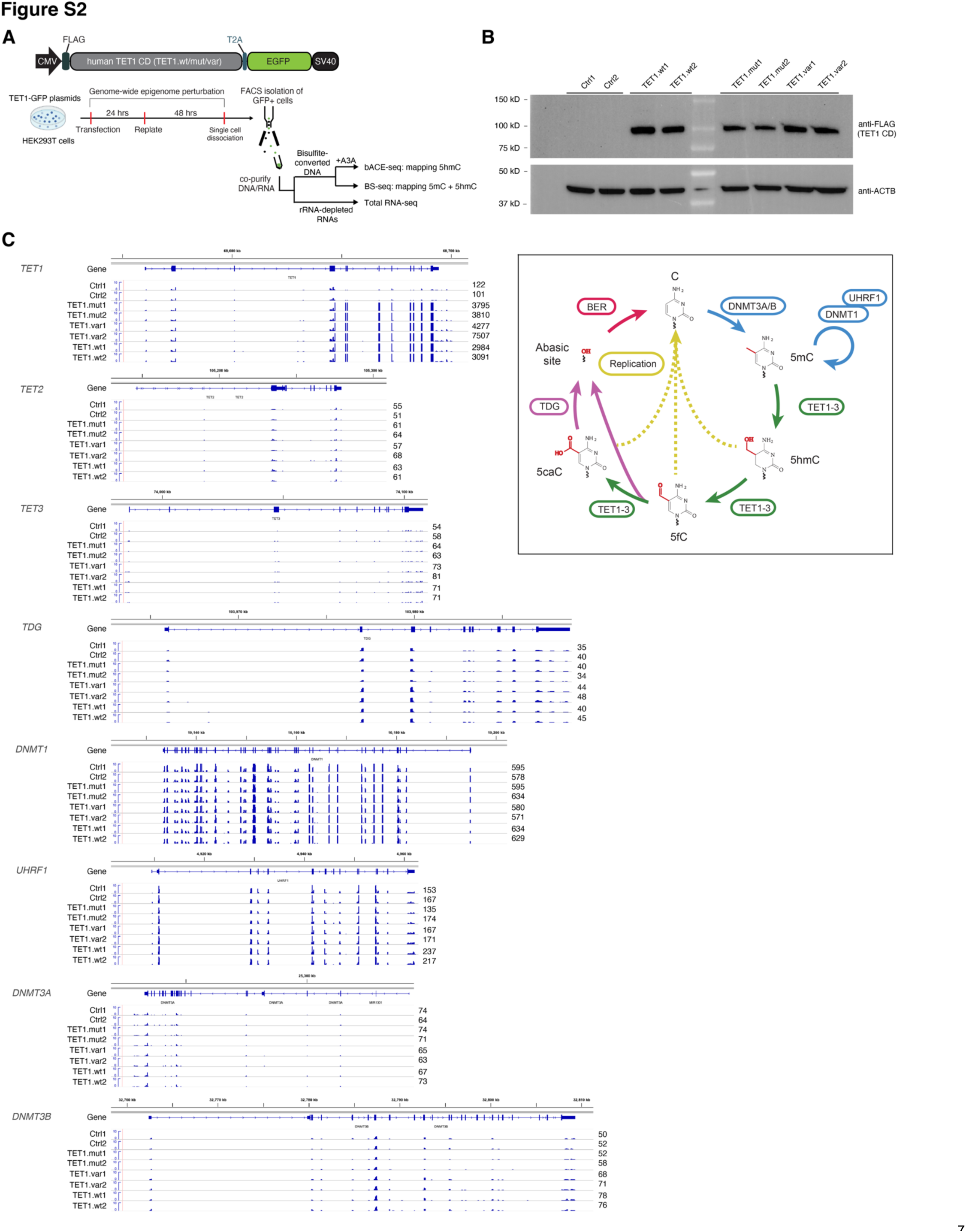
Experimental design and expression analysis for genome-wide epigenome editing in HEK293T cells. (**A**) Schematic experimental workflow for transfecting expression vectors encoding human TET1 (wt/mut/var) catalytic domains (CD) into HEK293T cells. Following transfection, cells are sorted with FACS to isolate GFP+/DAPI– cell populations. **(B)** Representative western blot analysis evaluating the protein expression of our recombinant plasmids in FACS-sorted GFP+/DAPI-cells. Top panel represents Flag-tagged human TET1.wt/mut/var CD; bottom panel denotes β-Actin, which is used as a loading control. Ctrl1, non-transfection control cells; Ctrl2, pTY-GFP expressing cells. **(C)** Genome browser view of gene annotation tracks and total RNA levels (scale of y-axis: 0 to 10) of genes encoding all major proteins involved in DNA methylation and demethylation dynamics in control (Ctrl1/2), TET1.mut, TET1.var, and TET1.wt expressing HEK293T cells, with normalized expression levels (in counts per million (CPM)) for each replicate shown next to the tracks. (Right) Diagram depicting the major components of the DNA methylation and demethylation cyclic cascade.

**Figure S3.**
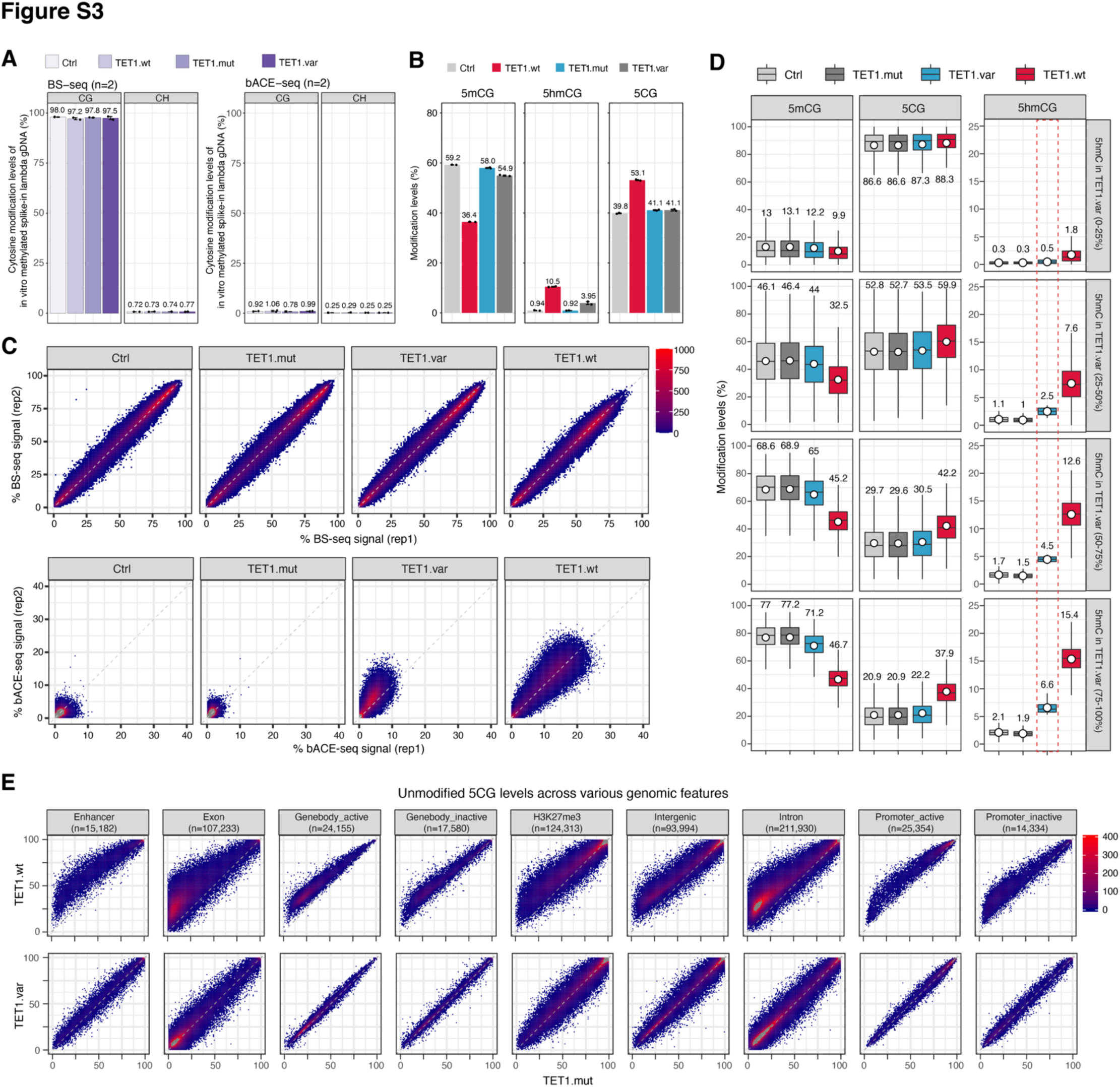
Validation experiments for integrated whole genome BS-seq and bACE-seq analyses. (**A**) Bar graph of 5mC levels within CpG and CpH contexts for *in vitro* methylated lambda phage genomic DNA (used as spiked-in controls in BS-seq and bACE-seq experiments), with 5mC levels (%) listed above each bar. **(B)** Bar graph of global levels of 5mCG, 5hmCG and 5CG in Ctrl and TET1.mut/var/wt-expressing cells, with cytosine modification levels (%) listed above each bar. **(C)** Correlation density plot between two replicates of BS-seq (top) or bACE-seq (bottom) for Ctrl and TET1.mut/var/wt-expressing cells. Correlation analysis is performed with all 10-kb genomic bins spanning the human genome. **(D)** Box plots of 5mCG, 5hmCG and 5CG levels at genomic regions enriched for different levels of 5hmCG in TET1.var-expressing cells (indicated by dashed red box). **(E)** Correlation density plot of 5CG levels between TET1.mut and TET1.wt (top) or TET.var (bottom) cells across all annotated genomic or regulatory regions, with the number of regions listed for each categroy.

**Figure S4.**
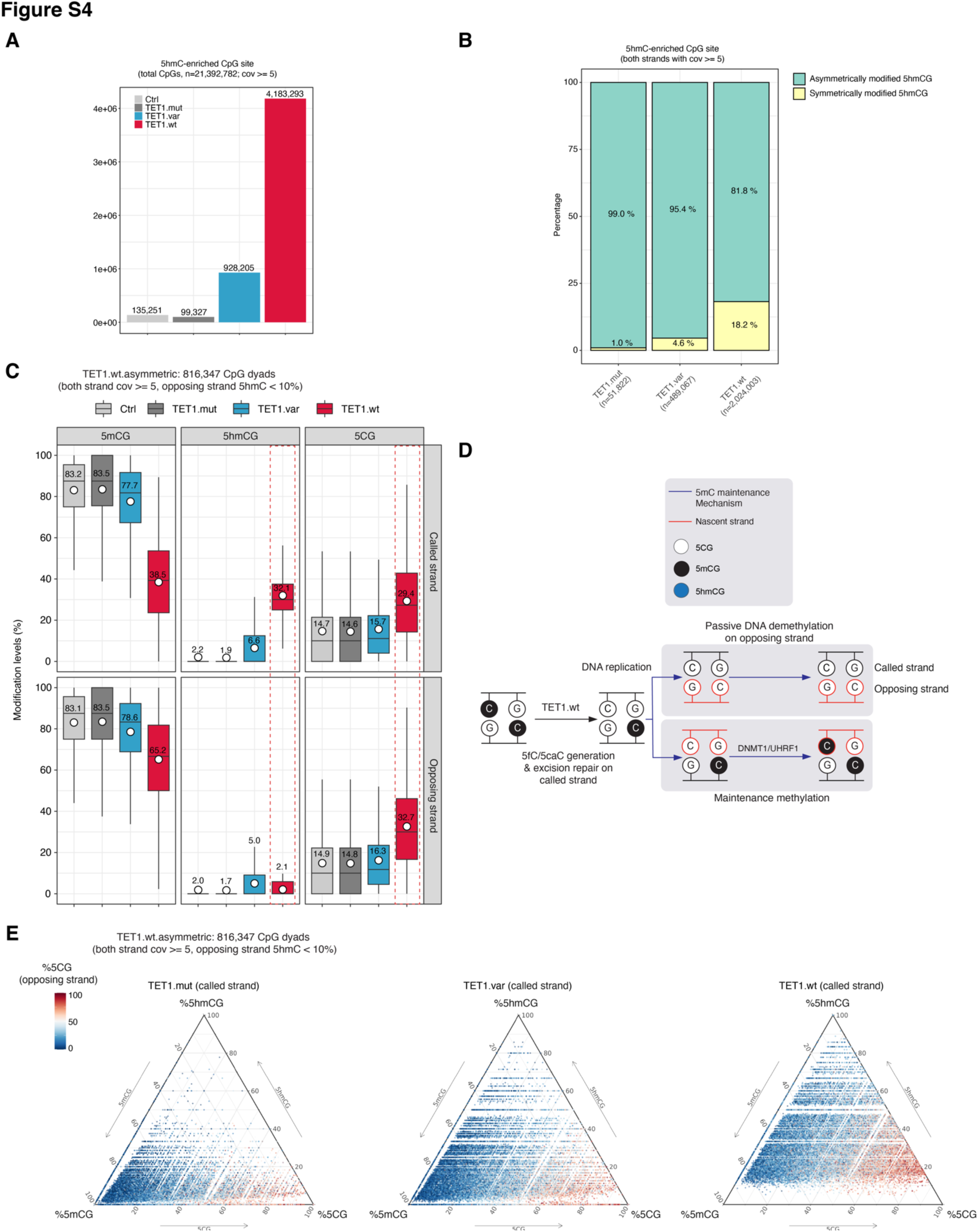
TET-mediated iterative 5mC oxidation is required for active DNA demethylation on both called and ing strands of CpG dyads in proliferating somatic cells. (**A**) Bar graph of CpG site (strand-specific) enriched for statistically significant 5hmC levels (*P* value = 2.5 x 10^-4^) in control and TET1.mut/var/wt-expressing human 293T cells, with the number of 5hmC-enriched CpG sites listed above each bar. **(B)** Bar graph showing the proportion of symmetrically (both strands modified) and asymmetrically (only one strand modified) hydroxymethylated CpG dyads in TET1.mut/var/wt-expressing HEK293T cells. **(C)** Box plots of 5mCG, 5hmCG and 5CG levels (%) on the called (top) and opposing (bottom) strands of asymmetrically hydroxymethylated CpG dyads (n = 816,347 sites) in TET1.wt expressing cells. **(D)** Schematic diagram of TET1.wt/TDG-mediated 5fC/5caC generation and excision repair and the impact of resulting hemi-methylation on the 5CG restoration on the opposing strand. **(E)** Ternary plots show the levels of 5CG, 5mCG and 5hmCG (%) on the called strand of asymmetrically hydroxymethylated CpG dyads (n = 816,347 sites) in TET1.wt expressing cells. TET.mut/var are shown as controls. The 5CG levels (%) on the opposing strand for the same CpG dyad are color coded (red: high; blue: low).

**Figure S5.**
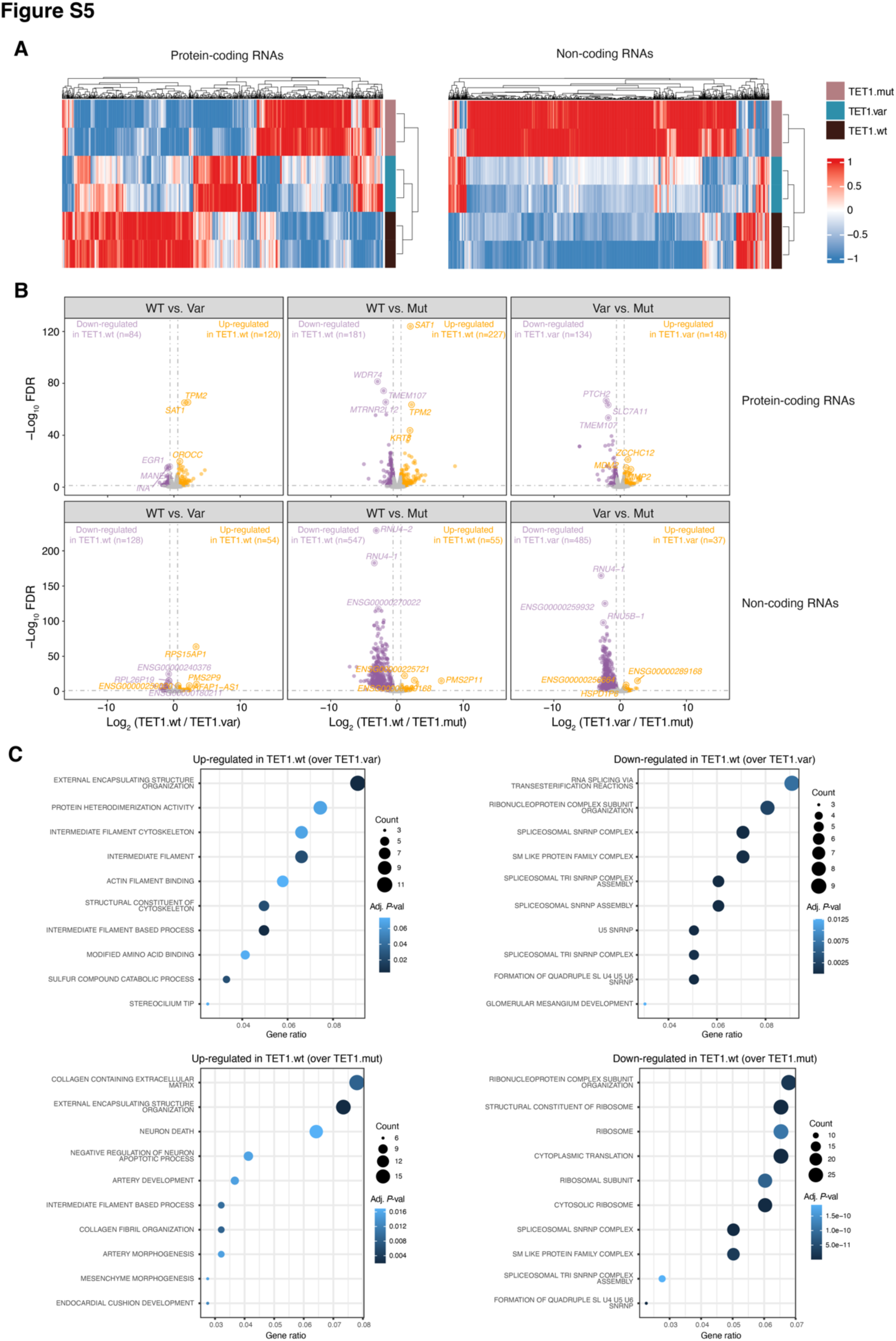
Total RNA sequencing analysis reveals the gene regulatory roles of TET-mediated stepwise, iterative xidation on protein-coding and non-coding RNAs. (**A**) Heat maps showing differentially expressed genes of protein-coding (left) and non-coding (right) RNAs among three pairwise comparisons: TET1.wt versus TET1.var (regulated by iterative 5mC oxidation alone), TET1.wt versus TET1.mut (regulated by iterative 5mC oxidation and/or 5hmC generation), and TET1.var versus TET1.mut (regulated by 5hmC generation alone). Both biological replicates are shown, and the relative gene expression is scaled by z-score. **(B)** Volcano plots of differentially expressed protein-coding (top) and non-coding (bottom) genes among three pairwise comparisons: TET1.wt versus TET1.var (regulated by iterative 5mC oxidation or 5hmC generation), TET1.wt versus TET1.mut (regulated by iterative 5mC oxidation alone), and TET1.var versus TET1.mut (regulated by 5hmC generation alone). Up-regulated genes are shown in orange, down-regulated genes are displayed in purple, and insignificant genes are shown in grey. **(C)** Top 10 enriched gene ontology terms for genes significantly up-regulated (left) or down-regulated (right) in pairwise comparisons [TET1.wt vs. TET1.var (top) or TET1.wt vs. TET1.mut (bottom)], respectively.

**Table S1.**
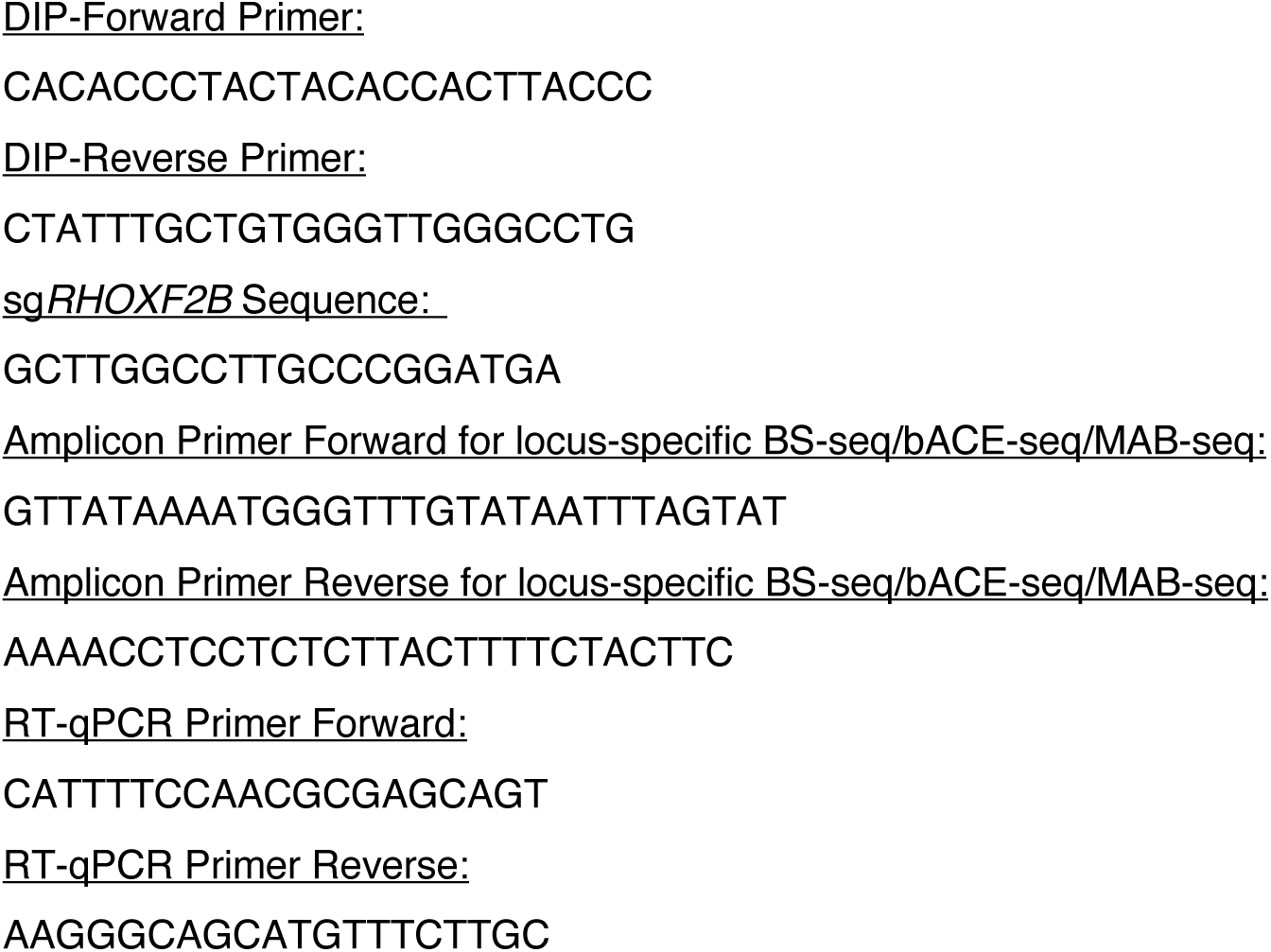
Oligonucleotides.

**Table S2.** Whole genome BS-seq and bACE-seq libraries.

| Samples | Method | Total reads | Uniquely mapped read | Mapping rate | Filtered reads (MAPQ>10) | Deduplicated reads | mCG_hg38 | mCHG_hg38 | mCHH_hg38 |
| --- | --- | --- | --- | --- | --- | --- | --- | --- | --- |
| Ctrl. rep1 | BS-seq | 334,288,342 | 238,498,927 | 71% | 211,530,779 | 193,058,053 | 60.40% | 1.05% | 0.90% |
| Ctrl. rep2 | BS-seq | 266,931,214 | 188,676,517 | 71% | 166,439,687 | 151,739,026 | 59.55% | 0.65% | 0.65% |
| TET1.mut, rep1 | BS-seq | 283,148,414 | 208,015,932 | 73% | 184,774,024 | 167,454,528 | 59.30% | 0.80% | 0.70% |
| TET1.mut, rep2 | BS-seq | 230,391,624 | 168,692,947 | 73% | 149,309,898 | 138,179,851 | 58.75% | 0.95% | 0.85% |
| TET1.var, rep1 | BS-seq | 369,668,236 | 262,061,271 | 71% | 230,813,407 | 211,071,722 | 58.40% | 1.00% | 0.90% |
| TET1.var, rep2 | BS-seq | 360,806,174 | 263,813,761 | 73% | 234,880,075 | 204,110,021 | 58.10% | 0.55% | 0.45% |
| TET1.wt, rep1 | BS-seq | 280,966,192 | 204,391,306 | 73% | 181,858,027 | 165,996,992 | 47.15% | 0.70% | 0.65% |
| TET1.wt, rep2 | BS-seq | 277,940,218 | 200,134,050 | 72% | 177,562,459 | 161,792,630 | 47.30% | 0.50% | 0.50% |
| Ctrl. rep1 | bACE-seq | 324,346,398 | 233,704,049 | 72% | 206,291,639 | 169,971,900 | 1.60% | 0.20% | 0.20% |
| Ctrl. rep2 | bACE-seq | 274,174,202 | 190,216,083 | 69% | 164,188,282 | 126,799,473 | 1.60% | 0.20% | 0.20% |
| TET1.mut, rep1 | bACE-seq | 387,222,684 | 291,336,125 | 75% | 259,149,190 | 207,691,499 | 1.30% | 0.20% | 0.20% |
| TET1.mut, rep2 | bACE-seq | 304,185,702 | 223,394,523 | 73% | 196,605,310 | 165,626,923 | 1.50% | 0.20% | 0.20% |
| TET1.var, rep1 | bACE-seq | 299,113,246 | 222,841,634 | 74% | 196,965,929 | 162,771,549 | 3.60% | 0.20% | 0.20% |
| TET1.var, rep2 | bACE-seq | 191,328,540 | 137,622,290 | 71% | 121,106,913 | 88,851,703 | 5.20% | 0.20% | 0.20% |
| TET1.wt, rep1 | bACE-seq | 239,583,876 | 178,930,525 | 74% | 158,917,149 | 126,722,086 | 10.70% | 0.20% | 0.20% |
| TET1.wt, rep2 | bACE-seq | 242,202,052 | 177,349,318 | 73% | 157,267,397 | 140,828,650 | 11.30% | 0.20% | 0.20% |

**Table S3.** RNA-seq libraries.

| Library type | Cell line | Samples | Total reads | Mapping rate |
| --- | --- | --- | --- | --- |
| Total RNA-seq | HEK293T | Ctrl, rep1 | 67,620,107 | 87.9% |
| Total RNA-seq | HEK293T | Ctrl, rep2 | 55,050,053 | 87.5% |
| Total RNA-seq | HEK293T | TET1.wt, rep1 | 63,936,554 | 89.1% |
| Total RNA-seq | HEK293T | TET1.wt, rep2 | 71,047,248 | 86.4% |
| Total RNA-seq | HEK293T | TET1.mut, rep1 | 78,802,333 | 80.5% |
| Total RNA-seq | HEK293T | TET1.mut, rep2 | 67,864,894 | 81.6% |
| Total RNA-seq | HEK293T | TET1.var, rep1 | 80,512,210 | 86.2% |
| Total RNA-seq | HEK293T | TET1.var, rep2 | 69,421,508 | 88.4% |

## Notes

### Competing Interest Statement

The authors have declared no competing interest.

